# Inferring gene regulatory networks from single-cell perturbation data by optimizing causal reach of latent mediator dynamics

**DOI:** 10.64898/2026.11.05.764481

**Authors:** Weizhou Qian, Yuxuan Song, Joshua D. Welch

**Affiliations:** Department of Computational Medicine and Bioinformatics, University of Michigan, Ann Arbor, MI, USA; Department of Computer Science and Engineering, University of Michigan, Ann Arbor, MI 48109, USA

**Keywords:** Causal Discovery, Gene Regulatory Networks, Perturbation, Ordinary Differential Equations, Machine Learning

## Abstract

Resolving the causal structure of gene regulatory networks is central to understanding cellular regulation. Pooled CRISPR screens with single-cell RNA readouts measure the transcriptional consequences of many interventions in parallel, providing a powerful resource for causal discovery. Combinatorial designs that perturb multiple regulators per cell increase throughput and can reveal responses that single-gene perturbations do not. However, inferring causal gene networks from these experiments remains challenging because gene regulatory interactions are nonlinear, dynamic, and often involve feedback. Thus, attributing the combined effect of several perturbations to individual regulators remains difficult. Additionally, transcriptional regulation is ultimately carried out by proteins, while single-cell perturbation datasets generally measure only RNA levels. Existing methods often simplify regulatory dynamics or rely on assumptions that conflict with gene regulation, such as acyclicity. Moreover, current evaluation practices usually rely on expression prediction error, which is readily computable in real datasets, but does not necessarily reflect graph quality. A complementary metric is needed to assess inferred regulatory connections using observed perturbation responses in real data.

To address these challenges, we present CausalODE, a differentiable framework for learning causal graphs from single-cell perturbation data. Our approach incorporates three central insights to improve upon previous methods. First, genes that are differentially expressed when a gene is perturbed should be downstream of the target gene in the regulatory network; we quantify this property with a metric we term causal reach. Second, causal interactions involve transient cellular responses in RNA transcription, splicing, and degradation. Third, gene regulatory interactions are mediated by proteins, whose levels are not directly measured in single-cell RNA-seq experiments. Building on these insights, CausalODE learns a graph that optimizes the causal reach metric, captures transient cellular responses to perturbation, and explicitly models the latent dynamics of protein levels. Together, these components allow nonlinear responses, feedback loops, and time-lagged regulation.

Across simulated Perturb-seq datasets with different sparsities and graph structures, CausalODE improves AUPRC and precision among top-ranked edges compared with baselines and competing methods, while achieving competitive AUROC. We introduce a metric to evaluate the causal reach of inferred graphs on real data by measuring how well the graphs explain differentially expressed genes in single-cell perturbation experiments. Using this metric, CausalODE outperforms the strongest competing method or baseline by approximately 60% on the CaRPool-seq knockdown dataset and 80% on the Norman et al. activation dataset. Its top-ranked edges generally show greater enrichment for high-confidence evidence from cell-type-matched ChIP-seq, CollecTRI, and regulatory STRING than those of other methods. The inferred graphs also distin-guish non-additive from approximately additive genetic interactions in held-out perturbation pairs. Finally, ablation analyses show that optimizing causal reach improves performance on held-out perturbations, and the learned latent regulator activities track simulated protein dynamics, with median temporal correlations ranging from 0.79 to 0.99.

## 1 Introduction

A gene regulatory network (GRN) describes how regulators control the expression of their target genes. Regulators often act through the molecules they encode: transcription factors bind regulatory DNA to activate or repress transcription, while signaling proteins alter downstream transcription factor activity. GRNs determine how changes in one gene propagate to others and shape cell fate over time. A better understanding of GRNs could create new possibilities for cell fate engineering and provide a more accurate blueprint for gene therapy[1, 2]. GRNs are also relevant to biological foundation models built from single-cell profiles[3]. However, these models often fail to generalize to unseen perturbations[4]. Several studies have argued that learning or embedding better GRNs could improve their generalizability[5, 6].

Methods such as GENIE3 infer regulatory relationships from statistical dependencies in observational expression data[7]. However, these dependencies may reflect direct regulation, indirect effects, or shared up-stream regulators. Pooled perturbation screens such as Perturb-seq measure single-cell responses to targeted gene intervention[8, 9]. CRISPR-based perturbations such as CRISPRi and CRISPRa can act as soft interventions on the GRN and, in some settings, as structural interventions, providing additional information that improves causal identifiability under appropriate assumptions[10, 11]. Many single-gene perturbations have weak effects because other regulators or redundant gene programs buffer their loss[8]. Combined perturbations can reveal these buffered responses and may enable causal graph identification with fewer experiments. However, testing all possible combinations is impractical.[12] Computational methods are therefore needed to attribute combined responses to individual regulators and infer causal GRNs from a limited set of perturbation combinations.

Under appropriate causal assumptions and study designs, a directed acyclic graph (DAG), or its Markov equivalence class, can be identified from data[13]. Constraint-based methods such as PC and FCI use conditional independence tests, which become computationally and statistically challenging for large GRNs[14, 15]. NOTEARS instead formulates DAG discovery as continuous optimization with a differentiable acyclicity constraint[16]. DCDI extends this approach to interventional data, while DCD-FG and SDCD improve scalability through restricted graph representations or sparse graph optimization[17, 18, 19]. However, DAG-based approaches cannot represent regulatory feedback loops. Dynamic models offer alternatives that allow cycles. BICYCLE models latent expression as the steady-state distribution of a linear stochastic dynamical system, limiting its ability to describe transient responses[13]. PerturbODE uses low-rank neural ODEs to model nonlinear cyclic dynamics[20]. A related dynamic method, RegVelo, models transcription and splicing through RNA velocity, but primarily refines a supplied GRN rather than discovering a causal graph de novo[21].

Graph evaluation presents a further challenge. Current evaluation often relies on expression prediction error or agreement with external references[13, 22]. However, accurate expression prediction does not necessarily imply a correct causal graph, because correlated expression programs can support predictions without the correct regulatory connections[23]. ChIP-seq and curated databases provide valuable supporting evidence, but physical binding or regulation reported in another context does not establish that an edge is causally active in the profiled cells. Complementary metrics are needed to assess inferred graphs directly using observed perturbation responses, including responses to unseen combinations.

Therefore, we present CausalODE, a differentiable framework for learning cyclic regulatory graphs by modeling RNA dynamics and delayed regulation through latent mediators while optimizing causal reach. The model combines transcription, splicing, and degradation with latent regulator activities and learnable Hill responses. Joint training with an RNA reconstruction loss and a DEG-based causal cross-entropy loss connects the modeled dynamics to significant perturbation responses and helps attribute combined responses to individual regulators. CausalODE supports knockdown and activation, single- and multigene perturbations, and single-snapshot or time-resolved data. We also developed causal reach violation (CRV) reduction, a new metric for evaluating inferred graphs when the true network is unknown. This metric measures how much better a graph connects perturbation targets to responding genes than randomly selected regulators, allowing paths of different lengths. To evaluate graph recovery and test the metric, we developed PerturbDyngen, an extension of dyngen that generates paired control and perturbed trajectories with matched stochastic sampling. We demonstrate improved graph recovery by evaluating CausalODE on simulated and real perturbation datasets. Additional analyses assess the causal cross-entropy loss, latent mediator dynamics, and biological support for the inferred graphs, and illustrate an application of the model.

## 2 Methods

We aim to infer regulatory graphs that explain significant perturbation responses while capturing RNA dynamics and delayed downstream regulation through unmeasured protein mediators. CausalODE couples RNA kinetics with latent mediator dynamics and uses causal reach supervision to constrain the inferred graph. The model takes per-cell unspliced and spliced counts, perturbation labels, and training perturbation DEGs as input. The dynamic model and causal loss share a weighted regulatory graph *W*, where *w*_*ji*_ describes regulation of gene *i* by regulator *j*. Candidate regulators *ℛ* are experimentally targeted genes, and *T*_*e*_ ⊆ *ℛ* denotes the targets of the perturbation group *e*. The following paragraphs describe the key components of CausalODE; further details are provided in **Supplementary Materials S2**.

### Transient RNA responses to perturbation

CausalODE models transcription, splicing, and spliced RNA degradation without requiring a steady state[24]:

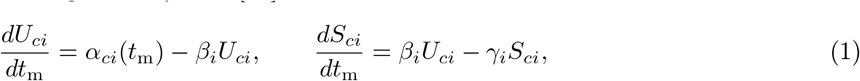

where *U*_*ci*_ and *S*_*ci*_ are the unspliced and spliced counts of gene *i* in cell *c*, and *t*_m_ is rescaled model time. The gene-specific splicing and degradation rates, *β*_*i*_ and *γ*_*i*_, are learned.

Perturbations scale target transcription by a measured multiplier *m*_*ci*_. Let *ρ*_*ci*_ be the target’s spliced expression divided by its mean in control cells, using all-gene library normalization before log transformation:

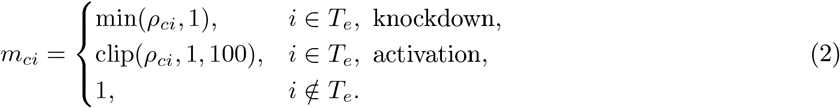

This retains only changes in the intended direction. For a zero control mean, we use complete knockdown or a reference activation multiplier equal to the larger of the mean and median activation multipliers of other targets in the training cells.

### Latent regulatory mediators

RNA changes affect downstream regulation through functional products such as transcription factor proteins. We represent their unmeasured activity by a latent relative activity

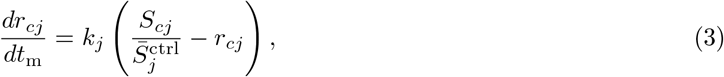

where 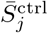 is the mean raw control spliced count. The learned relaxation rate *k*_*j*_ controls how quickly activity follows RNA changes, allowing delayed regulatory effects.

Latent activities control downstream transcription through Hill responses weighted by the graph:

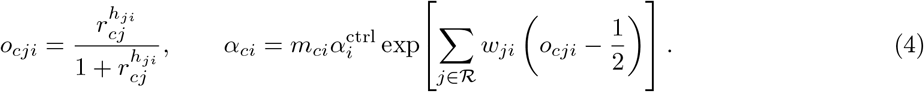

Here 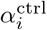 is a learned positive baseline transcription rate, and *h*_*ji*_ *>* 0 is a learned Hill coefficient that controls response steepness. The splicing, degradation, and relaxation rates (*β*_*i*_, *γ*_*i*_, and *k*_*j*_) are also learned and constrained to be positive. The Hill response depends on regulator activity and equals 1*/*2 at the control activity. Subtracting this control response makes each regulatory contribution reflect a deviation from the control baseline.

The graph, Hill coefficients, and kinetic parameters are shared across cells and perturbation groups. Effective transcription varies with perturbation strength and regulator activity. Feedback loops and autoregulation are allowed.

All cells start from mean raw control RNA counts with *r*_*cj*_ = 1. When sampling times are available, each cell is integrated to its sampling time. Single-snapshot data use a shared latent interval from *t*_0_ = 100 to *t* = 400. Because the full coupled system has no closed-form solution, we use numerical integration (Strang splitting) to solve the equations. The detailed algorithm is provided in **Supplementary Section S2.1**[25].

### Causal reach optimization and joint training

The causal cross-entropy loss constrains the graph to explain statistically significant perturbation responses. It encourages each DEG to be reachable from at least one perturbation target while discouraging links to non-DEGs.

We convert graph weights to edge probabilities and combine support from multiple targets:

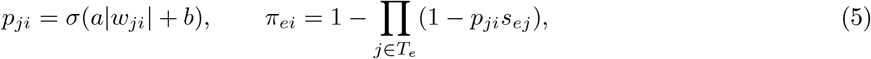

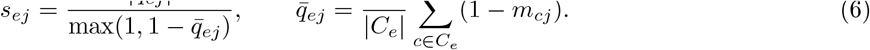

Where *a >* 0 and *b* are learned, *C*_*e*_ contains the cells in group *e*, and *s*_*ej*_ represents measured perturbation strength. A binary cross-entropy loss compares *π*_*ei*_ with DEG labels, giving DEGs and non-DEGs equal total weight. Target genes and self-edges remain included during training.

The graph and kinetic parameters are learned jointly by minimizing

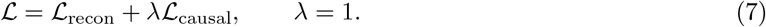

The reconstruction loss is the mean squared error between predicted and observed RNA counts after librarysize normalization over modeled genes and log(1 + *x*) transformation. Self-edges are removed at export because the perturbation design cannot reliably distinguish autoregulation from direct perturbation effects.

## 3 Results

### 3.1 CausalODE learns directed regulatory graphs from single-cell perturbation data

Our goal is to build a model that learns a weighted, cyclic directed regulatory graph by modeling RNA kinetics and latent protein mediator dynamics from single-cell perturbation data. We assume that perturbation-induced DEGs provide evidence of direct or indirect regulatory relationships and use these statistically significant responses to constrain graph learning. The model takes unspliced (*U*) and spliced (*S*) RNA counts, perturbation labels, and DEGs from each training perturbation group as input (**Fig. 1a**).

**Figure 1:**
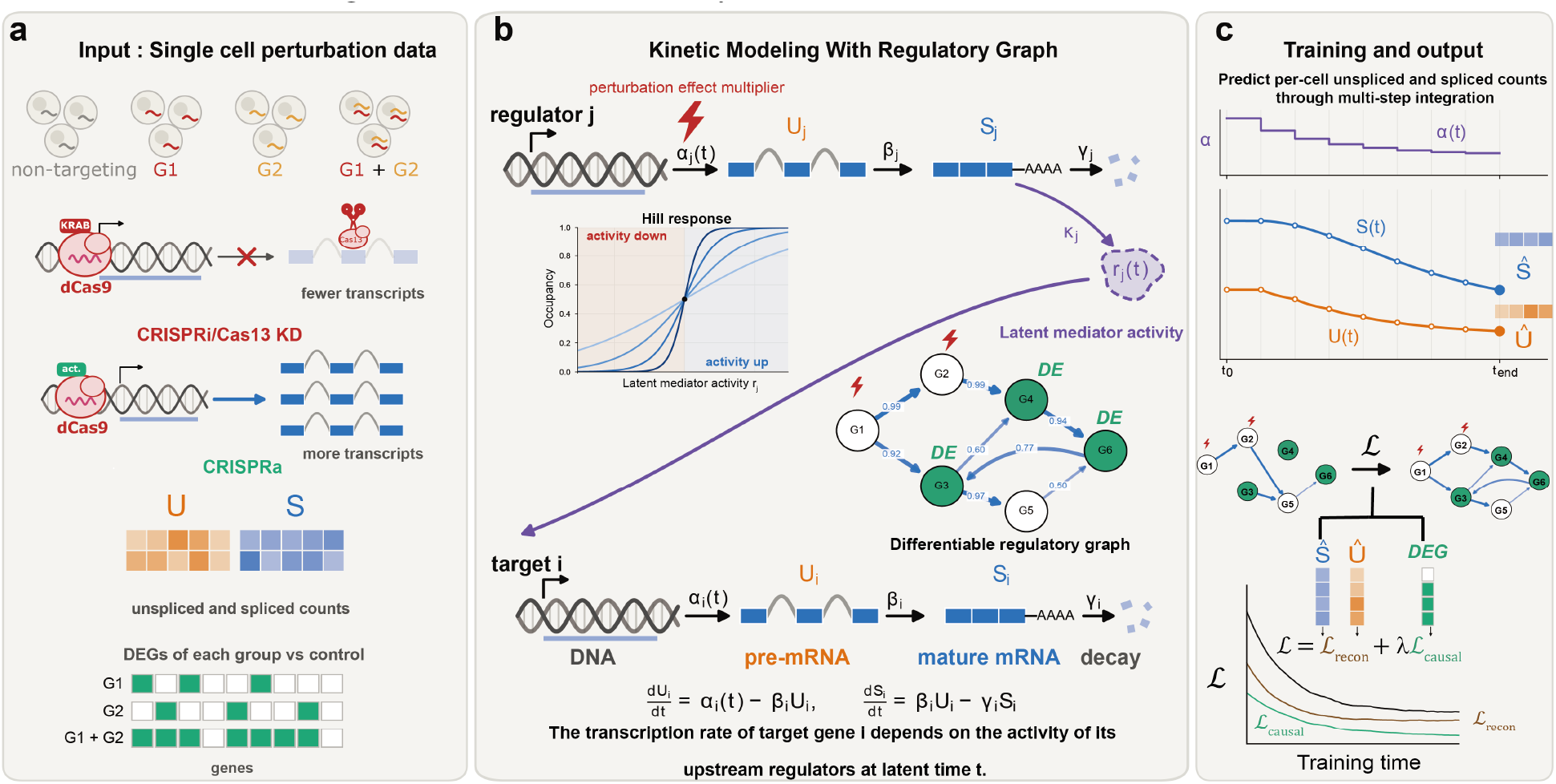
Overview of CausalODE. **(a)** CausalODE takes single-cell perturbation data (e.g., CRISPRi, Cas13, or CRISPRa) as input, with per-cell unspliced (*U*) and spliced (*S*) counts and per-group DEGs identified relative to non-targeting control cells. **(b)** RNA kinetics include transcription (*α*), splicing (*β*), and spliced RNA degradation (*γ*). Latent regulator activities control transcription through Hill responses weighted by a shared differentiable graph. Perturbations scale target transcription. **(c)** Multi-step integration predicts per-cell *U* and *S*, fitted to observations by *ℒ*_recon_. The causal loss encourages reachability of DEGs from at least one perturbation target and discourages reachability of non-DEGs. Graph and kinetic parameters jointly minimize *ℒ* = *ℒ*_recon_ + *λℒ*_causal_.

Starting from the mean state of control cells, our model uses kinetic ODEs commonly used in RNA velocity methods to simulate transcription, splicing, and spliced RNA degradation. Each gene has a time-dependent transcription rate *α*, a learned splicing rate *β*, and a learned degradation rate *γ*[24, 26]. To connect RNA expression with downstream regulation when regulatory mediators are unmeasured, we introduce a latent relative activity for each regulator (**Fig. 1b**). Although motivated by proteins encoded by regulatory genes, such as transcription factors, this variable *r* represents the effective activity of unmeasured regulatory products. Its dynamics approximate the production and turnover of these products without explicitly modeling translation or other molecular processes. The activity follows the regulator’s spliced RNA relative to its control level, allowing it to decrease or increase after knockdown or activation. A learned relaxation rate *k* controls how quickly the activity responds, allowing regulatory effects to lag behind changes in RNA expression.

Binding to regulatory sites can saturate, so changes in regulator activity need not produce proportional changes in target transcription. So learnable, edge-specific Hill functions (**Fig. 1b**) are integrated to the model to describe how regulator activity affects downstream transcription, capturing nonlinear responses with different steepness and saturation. Each response is centered at the regulator’s control activity, and the graph weights combine these responses to determine the target gene’s transcription rate relative to its control baseline. We assume that regulatory rules remain invariant within the training context. Therefore, graph weights, Hill parameters, and gene-specific baseline transcription, splicing, degradation, and activityrelaxation parameters are shared across cells and perturbation groups. The effective transcription rate *α* varies across cells and over time according to measured perturbation strengths and changing regulator activities.

The model allows feedback loops and autoregulation during training. When sampling times are available, each cell’s dynamics can be integrated to its sampling time. For a single snapshot, all cells share a latent integration interval without requiring the actual time elapsed since perturbation.

We train the graph and kinetic parameters jointly using two losses (**Fig. 1c**). The reconstruction loss matches predicted unspliced and spliced RNA counts to the observations, constraining the graph through the modeled dynamics. A DEG-based causal cross-entropy loss encourages links from perturbation targets to significant responding genes and discourages links to nonresponding genes. This compact, shared parameterization and the two simple losses support efficient training on larger gene panels. For each dataset, we select an early checkpoint after the reconstruction loss converges and improvements in validation causal loss become small. This helps prevent overfitting that can reduce graph quality. All training runs in this study met this rule (**Supplementary Fig. S1, S2**). Otherwise, we would select the checkpoint with the lowest validation causal loss. The final output is a weighted graph from the selected checkpoint, with self-edges removed because autoregulation cannot be reliably separated from direct perturbation effects.

### 3.2 CausalODE outperforms existing methods and baselines in simulated perturbation screens

To evaluate graph recovery against a known ground truth, we developed PerturbDyngen, an extension of dyngen for pooled perturbation screens (**Fig. 2a**)[27]. Given a regulatory graph and a cell-state transition map, the simulator generates single-cell trajectories through stochastic transcription, splicing, translation, and RNA and protein degradation. After a burn-in phase, each trajectory splits into paired control and perturbed continuations, with perturbations changing the transcription rates of selected genes. The two continuations use matched random sampling to control background stochastic variation. We then sample cells within a shared post-perturbation time window and add RNA capture noise to generate observed sparse unspliced and spliced counts.

**Figure 2:**
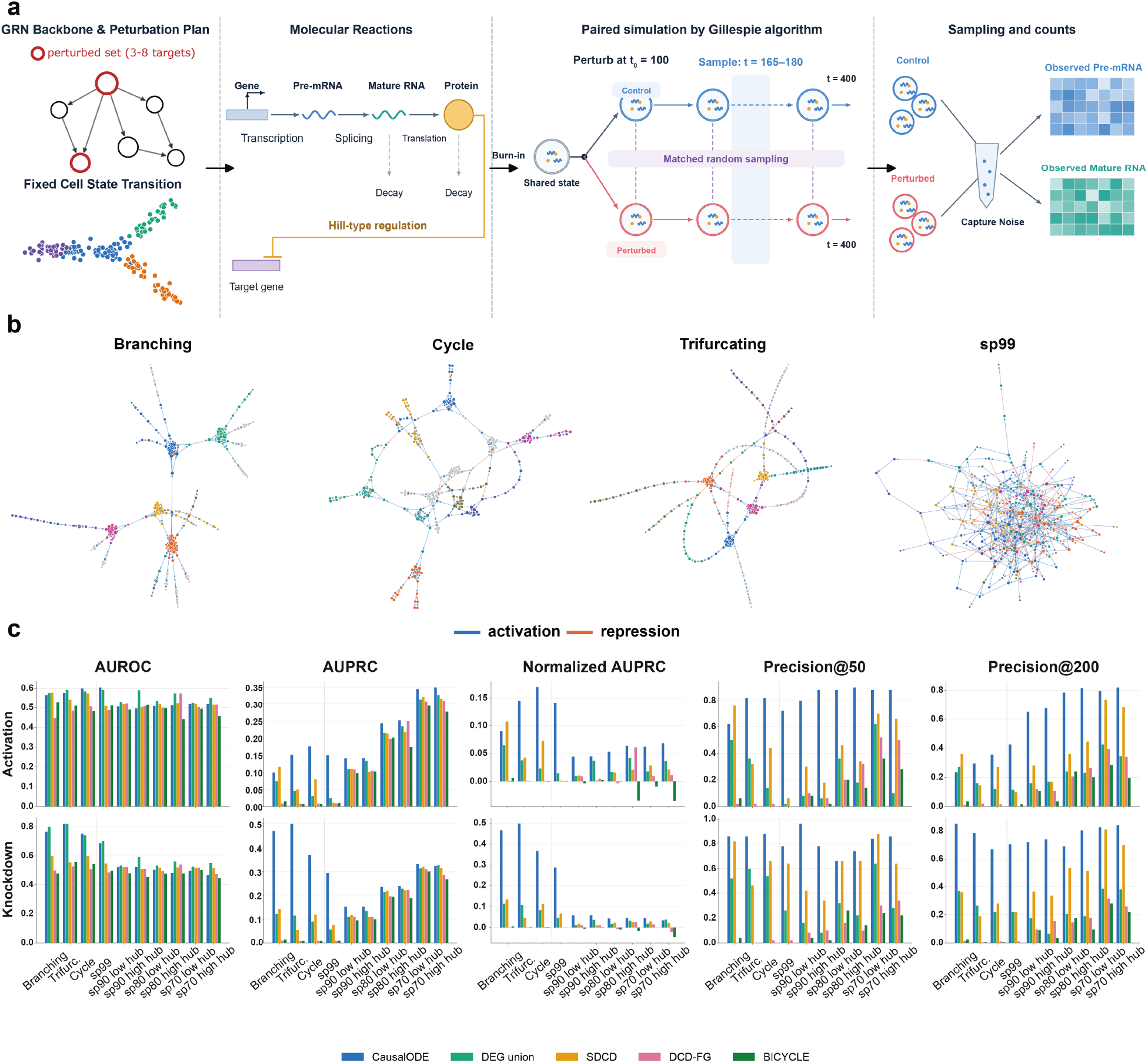
Causal GRN discovery benchmark using simulated single-cell perturbation datasets. **(a)** PerturbDyngen simulates RNA and protein dynamics in networks of 222 genes, with 76 perturbation conditions targeting 2–8 genes each. All regulatory network backbones share the same cell-state transition map. Paired control and perturbed trajectories use matched random sampling to avoid spurious differences caused by simulation randomness. Perturbations scale target transcription by 0.005 (knockdown) or 10 (activation) at *t* = 100. Cells are sampled at *t* = 165–180, followed by RNA capture noise. **(b)** Four example regulatory graphs. **(c)** Graph recovery across ten networks under activation (top) and knockdown (bottom), measured by AUROC, AUPRC, normalized AUPRC, and precision among the top 50 and 200 edges. The random networks sp99, sp90, sp80, and sp70 have 99%, 90%, 80%, and 70% sparsity, respectively. The latter three each have a low-hub and a high-hub version: in the high-hub version, a few TFs regulate many genes, while in the low-hub version, regulation is spread more evenly across TFs.

We simulated ten regulatory networks, each containing 222 genes, using the same cell-state transition map and kinetic parameter sampling distributions. **Fig. 2b** shows four example topologies. Three networks were derived from dyngen’s biologically motivated branching, trifurcating, and cyclic backbones. We also generated one random network with approximately 99% sparsity, matching the density of the branching backbone, and six denser random networks at 90%, 80%, and 70% sparsity, with low- and high-hub versions at each sparsity level. Each network was simulated with target transcription reduced to 0.5% of control for knockdown or increased 10-fold for activation, producing 20 datasets. Each dataset contained 76 combined-perturbation experiments (see **Supplementary Materials S3** for design details) targeting 2–8 genes each and 92,400 simulated cells. Statistics for all networks are provided in **Supplementary Table S1**.

We benchmarked CausalODE against SDCD, DCD-FG, BICYCLE, and the DEG union baseline[19, 18, 13]. DEG union constructs a graph directly from DEG labels, assigning each combined-perturbation response equally among the perturbed genes. This baseline tests whether CausalODE improves graph recovery beyond a direct attribution of DEG labels. We did not include PerturbODE because its implementation code was not publicly available at the time of benchmarking. Using the known regulatory graphs as ground truth and excluding self-loops, we evaluated AUROC, AUPRC, normalized AUPRC, and precision among the top 50 and 200 edges (**Fig. 2c**).

Across the 20 datasets, CausalODE achieved the highest AUROC or values close to the best-performing method. It achieved the highest AUPRC and normalized AUPRC on 18 datasets. The exceptions were the activation datasets for the branching network and the 70%-sparse, high-hub network, where CausalODE ranked second and performed close to the best method. CausalODE also recovered true edges with high precision among its top-ranked predictions, with a mean precision of 0.82 among the top 50 edges across datasets. It achieved the highest precision among the top 50 edges on 18 of 20 datasets and among the top 200 edges on 19 of 20 datasets. SDCD had slightly higher precision in the remaining cases.

Compared with DEG union, CausalODE achieved higher AUPRC and top-edge precision while maintaining comparable AUROC. These results suggest that CausalODE attributes combined perturbation responses to individual regulators more accurately than equal attribution of DEG labels. CausalODE generally performed better on knockdown datasets than on activation datasets. One possible explanation is that strong activation pushes Hill-type regulatory responses toward saturation, reducing their sensitivity to further changes in regulator abundance. Denser networks were also harder to recover. This may reflect greater regulatory redundancy: perturbing 2–8 genes leaves many other regulatory inputs intact, potentially buffering downstream responses.

### 3.3 CausalODE outperforms baselines and competing methods on unseen combined perturbations in real Perturb-seq datasets

We introduced causal reach violation (CRV) reduction to evaluate inferred graphs on real data because the true regulatory network is unknown (**Fig. 3a**). A CRV occurs when a differentially expressed gene cannot be reached from any perturbation target. Intuitively, a better graph should produce fewer CRVs for unseen combined perturbations. For each inferred graph, we compare violations for the actual perturbation targets with those for the same number of randomly selected regulators. To compare methods fairly, we convert edge rankings to probabilities at matched graph densities (0.1%–10%) and average the score across densities (**Supplementary Materials S4**). Because DEGs may respond indirectly to perturbations, we evaluate CRV reduction using paths of up to one, two, or three regulatory steps. Thus, CRV reduction has a direct interpretation: it measures how much better a graph with edge scores explains a held-out perturbation’s DEGs from its actual targets than from random regulators within the allowed number of steps, averaged across graphs constructed from those scores at matched densities. Positive values indicate better reachability than random regulators, zero indicates no improvement, and negative values indicate worse reachability.

**Figure 3:**
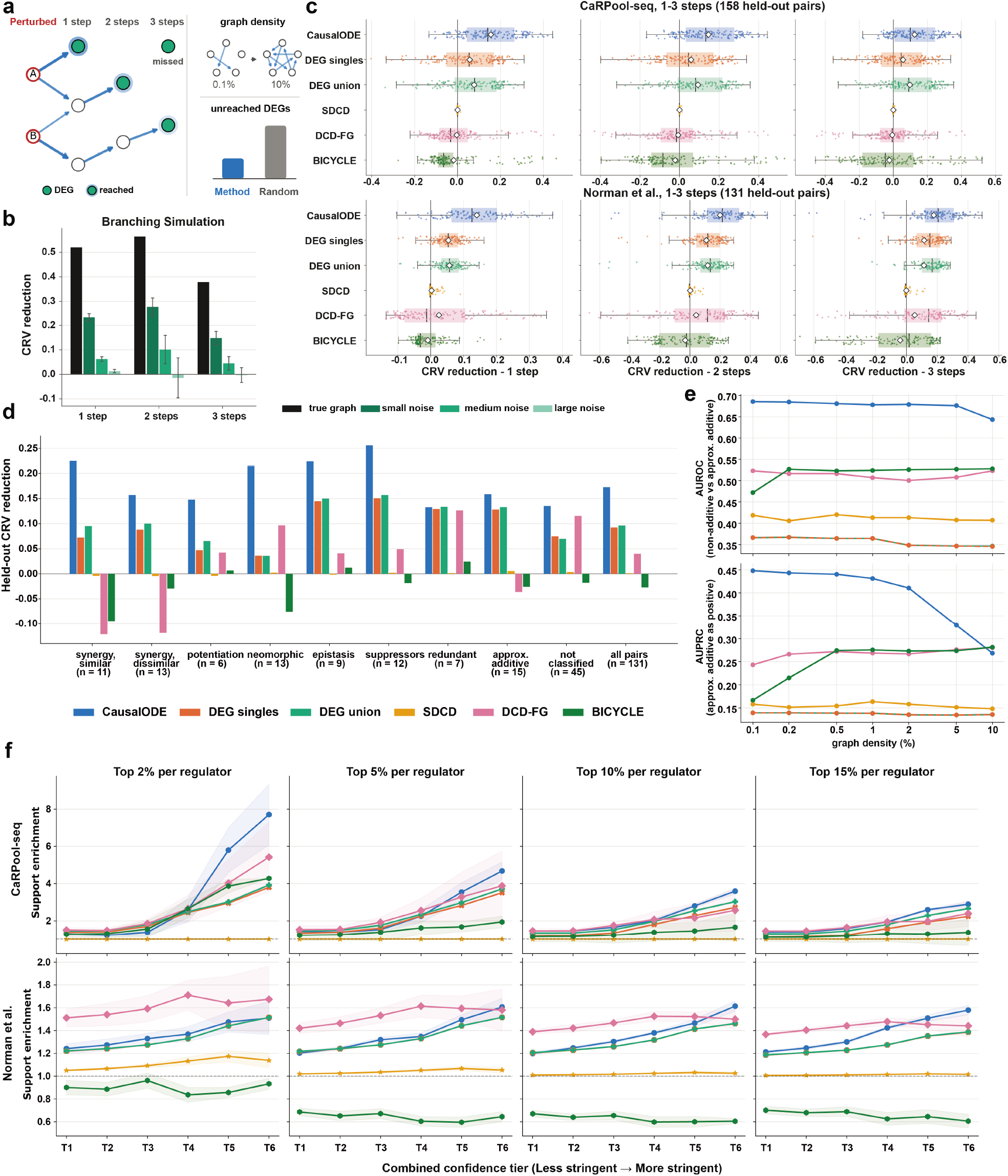
Evaluating inferred graphs with real perturbation data. **(a)** CRV reduction compares DEG reachability from perturbation targets with that from random stand-in regulators across matched graph densities (0.1%–10%). Higher scores indicate better graph quality. **(b)** Branching network: true graph and graphs with zero-mean Gaussian noise added to every gene pair (*σ* = 0.5, 1, 2), evaluated with one–three reach steps. Bars and error bars show means and s.d. over ten noise draws. **(c)** Held-out combined perturbations across five splits. Rows: CaRPool-seq and Norman et al.; columns: one-, two-, and three-step reach. Points represent pairs; diamonds show pooled scores. **(d)** Norman et al. CRV reduction by genetic-interaction type, averaged over one–three reach steps and graph densities. **(e)** Classification of 71 non-additive and 15 approximately additive Norman et al. pairs using co-reach. Each pair uses its corresponding split’s graph, trained without that pair. Top: AUROC for non-additive pairs. Bottom: AUPRC for approximately additive pairs, using negative co-reach. DEG singles and DEG union overlap. **(f)** External support enrichment for each regulator’s top 2%, 5%, 10%, and 15% of edges using matched-cell-type ChIP-Atlas, CollecTRI, or regulatory STRING evidence. Confidence tiers T1–T6 increase in stringency (**Supplementary Table S4**). Enrichment is relative to all evaluable edges from the same regulators. Lines and shading show means and s.d. across five splits.

We first tested whether CRV reduction reflects graph quality using the ten simulated networks and their knockdown datasets. We compared each true graph with graphs containing increasing Gaussian noise or other types of errors, such as extra false edges and shortcut edges. For each setting involving random corruption or random graph generation, we sampled ten graphs. Because these graphs were constructed rather than fitted, we used all perturbation experiments for this analysis. We focused on continuous noise because the methods in our benchmark also produce continuous edge scores.

In the Branching simulation, CRV reduction decreased monotonically as noise increased for one-, two-, and three-step reach (**Fig. 3b**). The true graph scored highest, followed by increasingly noisy graphs. This pattern held in all ten networks for one-step reach and at all three reach lengths in six networks (**Supplementary Fig. S3a**). In the other four dense networks (sp90 high hub, sp80 low and high hub, and sp70 high hub), adding small noise (*σ* = 0.5) produced slightly higher scores than the true graph at two steps, and also at three steps for sp90 high hub. However, scores still decreased as the noise level increased. Across networks, the median CRV reduction relative to the true graph decreased monotonically with increasing noise at every reach length. CRV reduction was also sensitive to other types of graph error (**Supplementary Fig. S3b**). For each error type and reach length, corrupted graphs had lower median scores than the true graph. More errors also produced lower median scores, and random graphs scored close to zero. These results show that CRV reduction generally ranks graphs by quality and is sensitive to all types of errors tested. Although two- and three-step scores can be less reliable for distinguishing small differences in some dense networks, median scores across networks consistently decreased as graph errors increased at all three path lengths. We retain all three path lengths because graphs inferred by different methods can differ substantially in practice, so two- and three-step CRV reduction should also help distinguish their quality. Allowing longer paths also accommodates indirect responses without assuming that every DEG is a direct regulatory target.

We then evaluated the methods on two screens with different perturbation modalities. CaRPool-seq used Cas13d to knock down 28 genes individually and in 158 pairs in THP-1 cells[28]. After filtering and restricting the analysis to GMP cells, the dataset contained 26,042 cells and a 1,000-gene panel. The Norman et al. screen used CRISPRa to activate 105 genes individually and in 131 pairs in K562 cells[8]. Its final analysis set contained 90,284 cells and a 2,000-gene panel. For each screen, paired perturbations were divided into five folds, with each pair held out once. Models were trained using single perturbations and the remaining pairs, then evaluated on the held-out pairs using one-, two-, and three-step CRV reduction (**Fig. 3c**). Because both screens include single-gene perturbations, we also included DEG singles, a baseline that links each perturbed gene to the DEGs observed after its single-gene perturbation in the training data.

CausalODE achieved the highest pooled CRV reduction on both datasets at all three reach lengths (**Fig. 3c**). After pooling held-out pairs across the five folds and averaging scores across reach lengths, CausalODE scored 0.142 on CaRPool-seq and 0.173 on Norman et al., approximately 60% and 80% higher than the strongest competing method or baseline, DEG union (0.089 and 0.096), respectively. DEG singles scored 0.057 and 0.093, while SDCD, DCD-FG, and BICYCLE scored substantially lower (**Supplementary Table S2**). CausalODE also achieved the highest scores at individual matched graph densities, except at 10% density with three-step reach (**Supplementary Fig. S4a**). Together with our simulation results linking CRV reduction to graph quality, these results indicate that CausalODE infers higher-quality regulatory graphs than the baselines and competing methods.

Norman et al. also annotated gene pairs by genetic-interaction type. In approximately additive interactions, the combined response is close to the sum of the individual perturbation responses. Non-additive interactions can amplify or suppress expression changes, so genes may become or cease to be differentially expressed under combined perturbations. These changes are not captured by simply combining DEG labels from the individual perturbations. Averaged across the three reach lengths, CausalODE achieved the highest or comparable CRV reduction across interaction types (**Fig. 3d**). DEG baselines performed similarly to CausalODE for redundant and approximately additive pairs. For redundant pairs, the combined response resembles a response already produced by an individual perturbation, whereas approximately additive pairs largely sum the two individual responses. Both cases are closer to a simple additive combination of existing perturbation effects and are therefore more readily captured by the DEG attribution used by DEG singles and DEG union. Similar patterns were also observed in the CaRPool-seq dataset using our own genetic-interaction annotations (**Supplementary Fig. S4b, S5**).

We next asked whether the inferred graphs could distinguish non-additive from approximately additive genetic interactions. Intuitively, two genes may interact non-additively when their effects converge on the same downstream genes. We therefore defined causal co-reach as the overlap between the downstream genes reachable from two regulators (**Supplementary Materials S4**). Using the graph from the fold in which each pair was held out, we evaluated 71 non-additive and 15 approximately additive Norman et al. pairs (**Fig. 3e**). CausalODE achieved the highest AUROC across graph densities from 0.1% to 10%, with values remaining around 0.65–0.70 when higher co-reach was used to predict non-additive interactions. We also evaluated AUPRC with approximately additive pairs as the positive class because they were the minority. We used negative co-reach because genes with less overlap in their downstream programs were expected to combine additively. CausalODE achieved the highest AUPRC at densities below 10%; at 10%, its performance was comparable to BICYCLE and DCD-FG. Non-additive pairs also had higher co-reach than approximately additive pairs in CausalODE’s inferred graphs (**Supplementary Fig. S6**), consistent with the expectation that overlapping downstream programs are associated with non-additive interactions. Together, these results provide additional evidence for the quality of CausalODE’s inferred graphs, which can predict non-additive genetic interactions in unseen perturbation pairs by capturing shared downstream regulation. In comparison, the competing methods and DEG baselines do not perform as well as CausalODE.

To better validate the inferred graph from different perspectives, we assessed external support for the top-ranked edges using cell-type-matched ChIP-seq data from ChIP-Atlas, CollecTRI, and regulatory STRING evidence (**Fig. 3f**)[29, 30, 31]. Combining these sources provided complementary biological evidence and increased coverage to 80.5% of candidate edges in CaRPool-seq and 58.1% in Norman et al. (**Supplementary Table S3**). We evaluated each regulator’s top 2%, 5%, 10%, and 15% of edges across six confidence tiers with increasingly stringent evidence thresholds (**Supplementary Table S4**). Enrichment was defined as the support rate among top-ranked edges divided by the support rate among all evaluable edges from the same regulators. On CaRPool-seq, CausalODE achieved the highest enrichment at both high-confidence tiers, T5 and T6, for all four edge fractions. On Norman et al., CausalODE achieved the highest enrichment at T6 for the top 10% and 15% of edges. Its enrichment was comparable to DCD-FG for the top 5%, while DCD-FG performed better for the top 2%. These results provide additional biological support for the quality of CausalODE’s inferred graphs, particularly its top-ranked edges, which generally show greater enrichment for high-confidence external evidence than those of competing methods.

### 3.4 Causal reach supervision improves graph inference and learned dynamics help explain perturbation responses

We examined the contribution of the causal cross-entropy loss by retraining CausalODE without this loss across all five folds of CaRPool-seq and Norman et al. The ablated model was trained using only the RNA reconstruction loss. Removing the causal loss substantially reduced CRV reduction on CaRPool-seq at every graph density (**Supplementary Fig. S7a**). The decrease was smaller on Norman et al., but the ablated model still scored lower at every density (**Supplementary Fig. S7b**). These results show that optimizing causal reach is important for improving graph quality.

We next examined whether CausalODE recovered the underlying kinetic parameters in the 20 simulated datasets. After normalizing learned values by their within-dataset medians, the pooled Spearman correlation with the true values was 0.24 for the spliced RNA degradation rate *γ* and 0.53 for the splicing-to-degradation ratio *β/γ* (**Supplementary Fig. S8**). We evaluated this ratio rather than *β* alone because the simulator used the same splicing rate for every gene. We did not directly compare transcription rates because CausalODE models transcription as time-dependent. These results indicate a limited ability to recover the true kinetic rates from a single snapshot.

Despite this limited rate recovery, the learned latent activities closely tracked simulated protein dynamics. We integrated the trained models from the control state and compared each perturbed regulator’s activity with its true protein abundance over 30 time points. Both quantities were expressed relative to their corresponding unperturbed controls at each time point. One knocked-down regulator in the Branching simulation showed a temporal Pearson correlation of 0.89 (**Supplementary Fig. S7c**). Across experiment–regulator pairs with a true protein change, median correlations were 0.99 and 0.79 under activation and 0.89 and 0.90 under knockdown for the Branching and sp99 networks, respectively (**Supplementary Fig. S7d**). Thus, latent protein activity captured temporal patterns in relative protein abundance even though kinetic rates were not accurately recovered.

Finally, we applied CausalODE to interpret gene-knockdown responses in CaRPool-seq. The original screen targeted regulators whose depletion was expected to promote monocyte differentiation, measured by increased CD11b. CD11b is encoded by *ITGAM*. After retraining CausalODE on the full dataset, we compared target RNA reductions with inferred activity reductions and regulator-to-*ITGAM* edge probabilities, using CD11b as the phenotypic readout (**Supplementary Fig. S7e**). This analysis revealed four patterns. In Type I, near-complete reductions in measured target RNA were accompanied by similar or weaker reductions in latent activity, high *ITGAM* edge probabilities, and increased CD11b. In Type II, latent activity decreased more strongly than target RNA. For example, *EP300, RCOR1*, and *DCAF1* perturbations showed little or no target RNA reduction but significantly increased CD11b. Their reduced latent activities and high *ITGAM* edge probabilities provided a possible interpretation of responses that were not apparent from RNA-based estimates of knockdown strength. In Type III, *CD14* and *ATXN7L3* showed strong RNA reductions and activity reductions but low *ITGAM* edge probabilities and no significant CD11b change, consistent with their inclusion as control genes in the original study[28]. Type IV showed high edge probabilities without significant CD11b changes and remained unexplained. These examples illustrate how CausalODE’s inferred graph and latent activities can help interpret perturbation responses beyond target RNA reduction alone.

## 4 Discussion

We developed CausalODE to infer cyclic regulatory graphs by modeling the causal reach of RNA dynamics and delayed regulation through latent mediators. The model scales to large gene panels with practical training times and is consistently much faster than BICYCLE in our benchmarks (**Supplementary Fig. S9**). Though the results show the model’s great ability to improve the causal graph recovery from perturbation data and CRV reduction shows promise as a proxy for graph quality, several limitations need to be emphasized.

CRV reduction depends on the graph density range and allowed path length, and does not consider regulatory signs. Although GRNs are generally considered sparse, choosing an appropriate density range remains difficult. Both the causal cross-entropy loss and CRV evaluation rely on DEG calls, which may miss weak or poorly sampled responses and may include false positives. Because the model relies heavily on interventional data, autoregulation cannot be reliably separated from direct perturbation effects, and inferred edges from genes that were never perturbed may be less reliable. The model also assumes shared regulatory rules and a common initial control state, which may miss differences between cell states or regulatory contexts.

The dynamical parameters require similar caution. With a single snapshot and a latent integration interval, absolute kinetic rates cannot be separated from the assumed time scale, and recovery of simulated degradation rates was limited. Latent activities tracked simulated protein dynamics but should not be interpreted as absolute measured protein abundance or half-lives. The unexplained Type IV CD11b examples may reflect graph errors or biological processes not adequately represented by the model. In addition, splicing and mRNA degradation rates are assumed to remain unchanged after perturbation, which may not hold in all settings.

Still, CausalODE, PerturbDyngen, and CRV reduction together provide a framework for learning and evaluating regulatory graphs from combinatorial perturbation data. Future work could test CausalODE on time-resolved perturbation data and use joint RNA–protein measurements and protein half-life information to validate and constrain the latent dynamics. Allowing splicing and degradation rates to vary, together with context-specific regulatory rules, could better capture diverse perturbation responses. Broader perturbation coverage could extend inference to more regulators, while further testing across density ranges, path lengths, and DEG-calling settings could clarify the robustness of CRV reduction.

## 5 Code and data availability

CausalODE is available at https://github.com/welch-lab/causalode, and PerturbDyngen is available at https://github.com/welch-lab/PerturbDyngen. The raw data for the Norman et al. and CaRPool-seq datasets can be accessed through GEO under accession numbers GSE133344 and GSE213957, respectively.

## 6 Acknowledgements

This work was supported by NIH grants R01HG010883 and U01HG011952 to JDW.

## Supplementary Materials

### S1 Dataset and preprocessing

#### Norman et al

We used the K562 CRISPRa Perturb-seq screen of Norman et al. (GEO GSE133344), in which cells carry one or two sgRNAs activating 105 genes singly and in 131 pairs. Spliced and unspliced counts were obtained from a published re-alignment of the raw reads, generated by pseudoalignment with kallisto (v0.46.2) and bustools (v0.40.0), run through kb-python (v0.26.0) with the La Manno spliced/unspliced workflow on the GRCh38 reference. Cells were matched by barcode and GEM group to the filtered cell identities released by Norman et al. (111,445 cells). Following the peprocessing in our PerturbNet, we kept cells with mitochondrial fraction *<* 10% of spliced and unspliced counts and ≥200 detected genes, and genes detected in ≥5 cells. We retained cells classified as singlets by the guide-calling procedure of Norman et al., leaving 90,284 cells (9,630 control, 46,779 single and 33,875 paired-perturbation cells).We then normalized spliced counts to 10,000 counts per cell, using the library size over all genes, followed by a log(1 + *x*) transformation. Highly variable genes were selected with Scanpy, with mitochondrial and ribosomal genes excluded from selection. The panel included all 105 target genes and the top 1,895 highly variable genes. Non-target genes that were never differentially expressed in any single perturbation were then replaced by the next-ranked highly variable genes, giving a final panel of 2,000 genes in which every non-target gene responds to at least one single perturbation. For benchmarking, we created five test splits containing only combined perturbations, such that every pair was held out exactly once, giving 26–27 test pairs per split.

#### Wessels et al

We used the CaRPool-seq screen from Wessels et al. (GEO: GSE213957), in which THP-1 cells carried Cas13d crRNA arrays targeting 28 genes individually and in 158 pairs. Spliced and unspliced counts were generated by processing raw reads with kallisto (v0.48.0) and bustools (v0.41.0) through kb-python (v0.27.3), using the GRCh38 reference and 10x Genomics v3 chemistry. We matched cells by barcode to the published per-cell metadata and retained only cells with annotated perturbation identities. Cell-type labels were provided by the authors. Using Scanpy (v1.10.3), we removed cells with fewer than 100 detected genes and genes detected in fewer than 3 cells, leaving 30,707 cells. We then retained cells annotated as granulocyte–monocyte progenitor (GMP), the only cell state with sufficient control cells, leaving 26,042 cells: 336 controls, 1,869 single-perturbation cells, and 23,837 paired-perturbation cells. We normalized spliced counts to 10,000 counts per cell using library sizes calculated across all genes, followed by a log(1 + *x*) transformation. Highly variable genes were selected using Scanpy based only on control and single-perturbation cells, excluding mitochondrial and ribosomal genes. The final panel comprised all 28 target genes and the top 972 highly variable genes, totaling 1,000 genes. For benchmarking, we constructed five test splits containing only paired perturbations, with each pair held out in exactly one split and 31–32 pairs per split.

#### Differentially expressed gene selection

For each perturbation group, we compared its cells with all control cells. Spliced counts were library-size normalized and log(1 +*x*)-transformed. We then applied a two-sided Wilcoxon rank-sum test using Scanpy and adjusted *P*-values with the Benjamini–Hochberg method. Genes with adjusted *P <* 0.05 were called DEGs. For the simulated datasets, DEGs were called in the same way using perturbed and control cells from the same sampling time window. The only difference was that we excluded the experiment’s target genes when calculating library sizes. Because only 222 genes were measured, changes in target-gene expression after knockdown or activation could otherwise shift library sizes and affect the normalized expression of all other genes.

### S2 Additional methods for CausalODE

CausalODE uses a kinetic model to describe how changes in latent regulator activity drive changes in each gene’s unspliced and spliced mRNA levels over time after perturbation. The model is trained using a reconstruction loss and a causal cross-entropy loss to learn a shared regulatory graph. The reconstruction loss matches predicted mRNA levels to the observed data. The causal cross-entropy loss uses observed DEGs to encourage a direct regulatory link from at least one perturbation target to each DEG, while discouraging such links to genes that do not respond.

The shared weighted regulatory graph is denoted by *W*, where *w*_*ji*_ represents the direct regulation of gene *i* by regulator *j*. Candidate regulators *ℛ* are genes targeted in at least one perturbation group. For perturbation group *e, T*_*e*_ ⊆ *ℛ* denotes the targeted genes.

The model requires three inputs: single-cell unspliced and spliced mRNA counts, each cell’s perturbation label, and the DEGs for each training perturbation group. Cell sampling times are optional. When cells are collected at multiple time points, each cell’s dynamics are integrated to its sampling time. For a single snapshot, as in all datasets in this study, all cells share a fixed latent interval after perturbation, from *t*_0_ = 100 to *t* = 400. The model learns relative changes in gene activity over this latent interval without requiring the actual time elapsed since perturbation.

Perturbations modify the transcription rates of targeted genes. We measure perturbation strength from each cell’s target expression. Spliced counts are normalized using the all-gene library size, and the expression ratio is calculated before log transformation. Let *ρ*_*ci*_ be the ratio of target *i*’s normalized spliced expression in cell *c* to its mean in control cells. We define

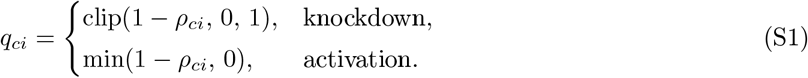

Only expression changes in the intended direction are retained; changes in the opposite direction receive *q*_*ci*_ = 0. If a target has zero mean expression in control cells, *ρ*_*ci*_ is undefined. For knockdown, we set *q*_*ci*_ = 1. For activation, we use the larger of the mean and median activation effects measured for other targets in the training cells.

For cell *c* in perturbation group *e*, the transcription-rate multiplier is

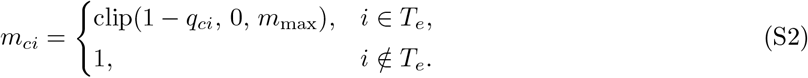

where *m*_max_ = 1 for knockdown and *m*_max_ = 100 for activation. Positive *q*_*ci*_ reduces transcription, whereas negative *q*_*ci*_ increases it. The lower bound of zero represents complete suppression of transcription.

#### RNA kinetics

For each gene *i* in cell *c*, unspliced mRNA *U*_*ci*_ is produced by transcription and converted to spliced mRNA *S*_*ci*_ by splicing. Spliced mRNA is then degraded. We write the dynamics in rescaled model time *t*_m_, defined in the numerical integration section:

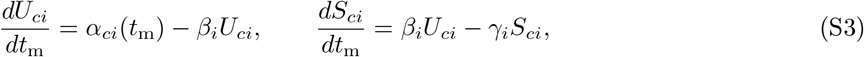

where *α*_*ci*_(*t*_m_) is the transcription rate, *β*_*i*_ is the splicing rate, and *γ*_*i*_ is the degradation rate of spliced mRNA. The splicing and degradation rates are learned for each gene and kept positive through softplus transformations:

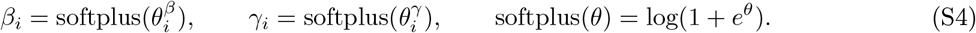

The parameters 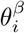 and 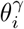 are unconstrained real numbers.

#### Latent transcriptional regulation

Regulatory genes act through their functional products, such as transcription factor proteins. Because the activity of these products is not directly measured, we represent each regulator *j* in cell *c* by a latent relative activity *r*_*cj*_(*t*_m_). This activity follows the regulator’s spliced mRNA relative to its control level:

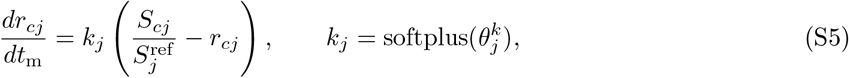

where 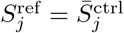 is the mean raw spliced count of regulator *j* in control cells and *k*_*j*_ is a learned relaxation rate. This allows changes in regulatory activity to lag behind changes in mRNA, with larger *k*_*j*_ giving a faster response. The parameter 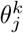 is learned without constraints.

The transcription rate of gene *i* in cell *c* is

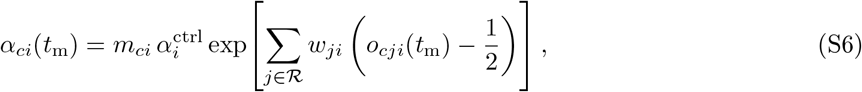

where *m*_*ci*_ is the perturbation multiplier and 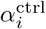 is a learned baseline transcription rate. The sum includes *j* = *i* when *i* ∈ *ℛ*, allowing autoregulation during training. In this model, perturbations and changes in regulator activity affect transcription only; each gene’s splicing and degradation rates remain constant across time, cells, and perturbation groups. Each regulator contributes a Hill response centered at its control baseline and weighted by the learned regulatory edge weight. The model assumes that these contributions are summed inside the exponential to determine their combined effect on transcription.

The regulatory response *o*_*cji*_(*t*_m_) is a Hill function of the relative activity:

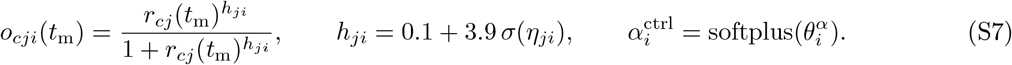

Here *σ* is the logistic function, *h*_*ji*_ ∈ (0.1, 4) controls how gradually or sharply the target responds to changes in regulator activity, and *w*_*ji*_ is the regulatory weight. The parameters 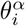, and *w*_*ji*_ are learned without constraints.

For each regulatory edge, we place the half-maximal Hill response at the regulator’s control activity: when *r*_*cj*_ = 1, *o*_*cji*_ = 1*/*2. This puts the control activity in the middle of the response range, allowing both increases and decreases in regulator activity to affect target transcription. Subtracting 1*/*2 centers each regulatory contribution on this reference, so the transcription rate is equal to 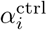 when all regulators are in their control activities and *m*_*ci*_ = 1.

The same regulatory functions, weights, and Hill coefficients are used for all cells and perturbation groups. Cells respond differently through their perturbation multipliers and the resulting regulator activities, while the underlying regulatory rules remain shared.

### S2.1 Numerical integration

We integrate the coupled dynamics using fixed-step operator splitting, with an exact solution for each sub-step. Sampling time *t* is converted to model time by

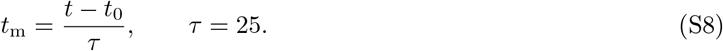

The rescaled interval is divided into *N* = 8 equal steps of length

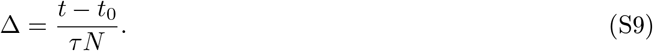

For *t*_0_ = 100 and *t* = 400, integration runs from *t*_m_ = 0 to *t*_m_ = 12, with Δ = 1.5. The time scale *τ* fixes the units of the learned kinetic rates. When cell-specific sampling times are available, each cell uses its own integration interval.

Steps are indexed by *n* = 0, …, *N* − 1, with step *n* covering [*n*Δ, (*n* + 1)Δ] in model time. All cells start from the control means at *t*_m_ = 0, corresponding to *t* = *t*_0_:

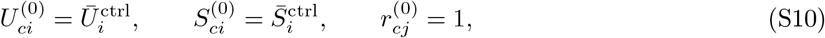

where 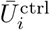 and 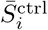 are the mean raw unspliced and spliced counts of gene *i* in control cells. The activity reference is 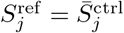. Each step updates the state in three parts:

1. *First activity half-step*. Spliced counts are held fixed at 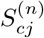, and regulator activities are updated over

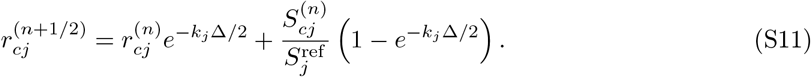

Here 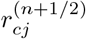 denotes the intermediate activity after this half-step.
2. *Full RNA step*. The transcription rate 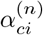 is computed from the intermediate activities 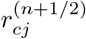 using the regulatory model above and held constant over Δ. With this fixed transcription rate, the RNA equations are linear and have an exact solution. Define

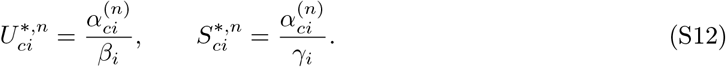

These are the levels that unspliced and spliced RNA would approach if the current transcription rate remained constant. They are recomputed at each step. The RNA update is

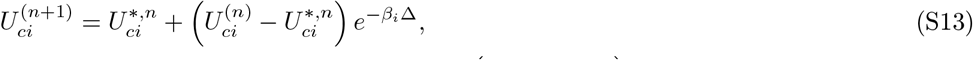

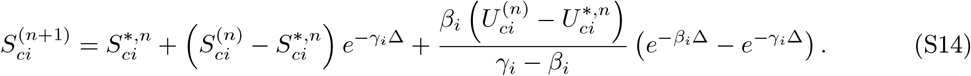
3. *Second activity half-step*. Regulator activities are updated over another Δ*/*2, now using the updated spliced counts:

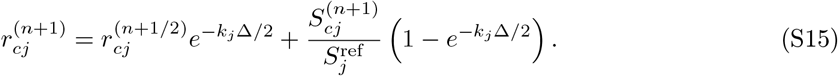

Each sub-step is solved exactly, while the coupled dynamics are approximated by alternating the activity and RNA updates. After *N* steps, the predicted counts 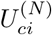 and 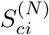 are used in the reconstruction loss.

#### Training objective

CausalODE is trained by minimizing

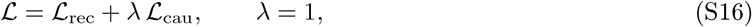

where *ℒ*_rec_ is a reconstruction loss and *ℒ*_cau_ is the causal cross-entropy loss. Both are computed on minibatches of cells.

##### Reconstruction loss

For each cell, the kinetic model predicts unspliced and spliced counts *Û*_*ci*_ and *Ŝ*_*ci*_. Predicted and observed counts are normalized by library size over the modeled genes *G* and transformed using log(1 + *x*), yielding 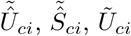, and 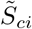. For a mini-batch *B*, the reconstruction loss is

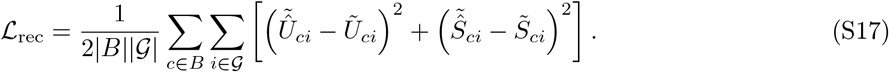

Control cells contribute only to this loss.

##### Causal cross-entropy loss

The causal loss encourages the graph to explain which genes respond to each perturbation. The magnitude of each regulatory weight is mapped to an edge probability using a learned linear transformation followed by a sigmoid function:

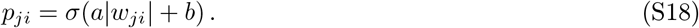

For this loss, the strength of target *j* in perturbation group *e* is averaged over the group’s cells *C*_*e*_:

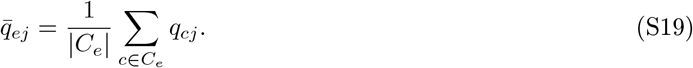

The predicted probability that gene *i* responds is a noisy-OR over the group’s targets:

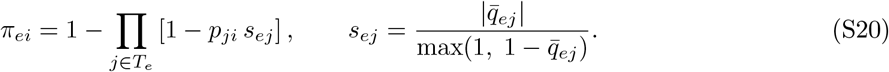

This combines support from individual targets, with stronger edge probabilities and stronger perturbations increasing the predicted response probability. Here 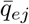 is the mean perturbation effect of target *j* over the cells of group *e*, with each cell’s effect capped at the same bound as in the ODE (*m* ≤ *m*_max_ = 100, i.e. *q* ≥ 1 − *m*_max_). For knockdown 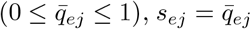, the fraction of transcription removed. For activation (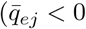, transcription multiplied by 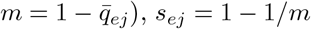), *s*_*ej*_ = 1 1*/m*.

Let *D*_*e*_ be the observed DEGs of group *e* and 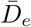 the remaining modeled genes. Target genes are not excluded from these labels, and diagonal probabilities *p*_*jj*_ are retained in the noisy-OR. Thus, a target’s own response can be supported by its self-edge as well as by edges from other targets in the group.

Because DEGs usually form a small fraction of genes, DEGs and non-DEGs receive equal total weight:

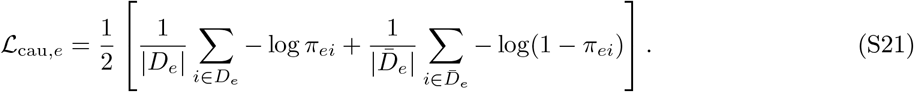

The overall causal loss _cau_ will train the graph to explain the DEG patterns across all training perturbations, including combinations of targets.

### S2.2 Optimization and graph selection

All parameters are optimized jointly with Adam on mini-batches of 128 cells, with gradient-norm clipping at 2. The kinetic parameters use a learning rate of 4 × 10^−3^. The graph weights *w*_*ji*_ and calibration parameters *θ*^*a*^ and *b* use a smaller learning rate of 4 × 10^−5^ to update the graph more gradually. Weights are initialized at *w*_*ji*_ = 0.05 and Hill coefficients at *h*_*ji*_ = 1.

About 10% of the training perturbation groups, drawn at random from the combinations, were held out as validation groups for checkpoint selection. Training runs for up to 120 epochs. We select an early checkpoint after the reconstruction loss has converged and improvements in the validation causal loss have become small. Starting at epoch 10, the three-epoch moving average of each validation loss is compared with its value five epochs earlier, and a loss is considered converged when its relative improvement is below 3%. The selected epoch is the first at which both losses meet this condition for three consecutive epochs. This rule is intended to limit overfitting of the graph once expression reconstruction has converged and further graph improvements are small. If no epoch meets it, the epoch with the lowest validation causal loss is selected; all training runs in this study met the rule (**Supplementary Fig. S1, S2**).

The edge probabilities *p*_*ji*_ at the selected epoch form the exported graph, with self-edge scores *p*_*jj*_ set to zero. Diagonal weights and probabilities remain trainable throughout fitting and are masked only during export. Because the current perturbation design does not reliably distinguish autoregulation from the direct effect of perturbing a target, we do not interpret self-edge scores as evidence of autoregulation.

### S3 Simulation

We developed PerturbDyngen, a simulator of pooled perturbation screens built on dyngen. In dyngen, each gene is represented by pre-mRNA, mature mRNA, and protein, linked by transcription, splicing, mRNA decay, translation, and protein decay reactions. Transcription rates depend on regulator protein levels through Hill-type activation and repression functions. The resulting chemical reaction network is simulated using GillespieSSA2 with tau-leaping, which advances the system in discrete time steps by sampling the number of events for each reaction from a Poisson distribution. Each simulated trajectory represents one cell. We used networks of 222 nodes. Our main extensions were to synchronize the random sampling of paired control and perturbed trajectories and to simulate noise in guide detection and counts. We also implemented a custom sequencing-noise model.

Each trajectory was first simulated with control parameters through a burn-in phase and until the perturbation time *t* = 100. From this shared state, we simulated two continuations until *t* = 400: one with control parameters and one with the transcription rates of targeted genes multiplied by a perturbation strength, below 1 for knockdown and above 1 for activation. In dyngen’s tau-leaping implementation, Poisson sampling can consume different numbers of random draws depending on the reaction rate. Consequently, using the same random seed does not keep the two trajectories synchronized after perturbation, potentially introducing apparent expression differences even in genes unaffected by the perturbation. To maintain synchrony, we used common random numbers: at each shared time step, we generated one uniform random number per reaction and used it to sample event counts for both trajectories by inverse-CDF Poisson sampling. Reactions with unchanged rates therefore received identical event counts, ensuring that differences in simulated molecule counts between paired control and perturbed trajectories arose from the perturbation rather than mismatched random sampling.

To generate observed RNA counts, we applied sequencing noise to the true molecule counts at each snapshot. Each cell *c* was assigned a size factor *s*_*c*_ ~ LogNormal(0, 0.1). For each gene *g*, we sampled an observed count from a negative binomial distribution with mean *s*_*c*_*pn*_*cg*_ and dispersion *θ* = 3, where *n*_*cg*_ is the true molecule count and *p* is the capture probability: 0.05 for mature mRNA and 0.01 for pre-mRNA. Sampled counts were capped at *n*_*cg*_.

To mimic guide measurement noise in pooled screens, PerturbDyngen also simulates guide UMI counts using a sigmoid relationship between guide abundance and perturbation strength, with negative binomial noise, cell-specific size factors, and incomplete detection. We omit implementation details because guide counts were not used in this study.

#### Benchmark graphs and experimental design

Because known regulatory graphs are sparse, we used three sparse graph backbones from dyngen, which have biologically motivated structures: branching, trifurcating and cycle. To test generalizability, we also sampled a random network with the same density as the branching backbone (99% sparsity), and denser random networks at 90%, 80% and 70% sparsity, each with a low and a high hub level. In the sampled networks, out-degrees followed a log-normal distribution, with its spread controlling the hub level. Each edge was initially assigned as activating or repressive with probability 0.75 or 0.25, respectively. So the expected number of repressive inputs per gene increased with network density. Genes with many repressive Hill inputs were permanently silenced in the simulations. To avoid this, we limited each gene to at most three repressive inputs by randomly changing excess repressive edges to activating edges. This reduced the overall fraction of repressive edges in denser networks. In total, we simulated 10 graphs (**Supplementary Table. S1**).

To test whether a small number of strong interventions makes the graph identifiable, each experiment perturbs 2 to 8 genes at once. For each graph, 57 experiments (7–8 genes each) were designed to satisfy the ordered-pair condition: for any two genes *i* and *j*, at least one experiment perturbs *i* but not *j*, and at least one perturbs *j* but not *i*. Another 19 experiments target the upstream regulators of the genes with the most inputs, so that their responses are not masked by the remaining regulators. Knockdown multiplied the transcription rates of the targeted genes by 0.005, and activation multiplied them by 10. We selected cells from a manually chosen time window, *t* = 165–180 (65–80 time units after perturbation), when perturbation responses were well developed and indirect effects were present but did not dominate as they did at later time points. Each final dataset contained 92,400 simulated cells.

#### Kinetic parameters and simulated cells

All networks used dyngen’s default kinetic sampling, drawn independently for each gene and network. Transcription rates were drawn uniformly from 10–20, translation rates from 100–150, mRNA half-lives from 2.5–5 and protein half-lives from 5–10, the latter multiplied by three to give 15–30 so that regulator proteins change more slowly than their mRNA. The splicing rate was the same for every gene (log 2*/*2 ≈ 0.35). Every regulatory edge had strength 1 and a Hill coefficient of 2 or 4, and regulatory inputs to each gene were combined with dyngen’s thermodynamic model. Basal transcription was 1 for the burn-in root gene and 0.2 for all other genes. Thus networks differ in topology and in the individual kinetic draws, but all draws come from the same distributions.

Trajectories start with no molecules, run through a burn-in of 180 time units with a tau-leaping step of 300*/*3600 time units, and are recorded every 5 time units. All networks use the differentiation path of the branching network as dyngen’s backbone definition. Every experiment uses the same 300 trajectory seeds, so the control continuations are identical across experiments.

### S4 Benchmark metrics

#### Simulation

We compared each method’s edge scores with the simulator’s graph, excluding self-loops, using the area under the ROC curve (AUROC), the area under the precision–recall curve (AUPRC), the normalized AUPRC, (AUPRC − *π*)*/*(1 − *π*) with *π* the edge prevalence, and the precision of the top-ranked edges, Precision 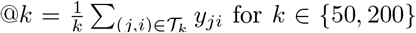, where *T*_*k*_ is the set of *k* gene pairs with the largest absolute scores and *y*_*ji*_ = 1 if *j* → *i* is an edge of the simulator’s graph.

#### Causal reach violation (CRV) reduction

Without a known graph in real data, we evaluated whether an inferred graph explains the responses to unseen perturbations. The DEGs of a perturbation must be reachable from the perturbed genes; a DEG that cannot be reached, for example one upstream of or in a parallel branch to the perturbed genes, is a causal reach violation. To compare methods with different score scales, we converted scores into density-matched edge probabilities. The absolute scores of all off-diagonal edges from perturbed genes, *N* = |*P*|(*n* − 1) entries, were ranked (*r*_*ij*_ = 1 for the strongest), and for *k* expected targets per regulator, *d* = *k/*(*n* − 1),

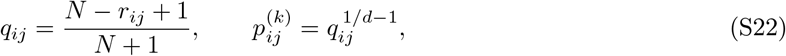

with tied scores sharing their mean probability. For a held-out pair *e* with perturbed genes *T*_*e*_ and DEGs *D*_*e*_ (significant non-target genes), reach within *L* steps is computed as a noisy-OR over paths through perturbed genes:

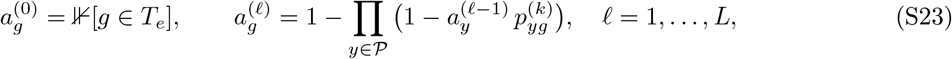

with perturbed genes kept at *a*_*g*_ = 1, and

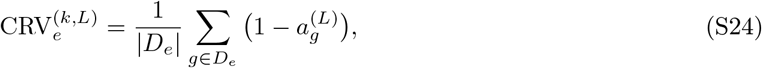

the probability that a DEG is not reached from the perturbed genes within *L* steps; for *L* = 1 this is 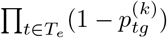. As a reference, we replaced *T*_*e*_ with the same number of randomly chosen other perturbed genes (*B* = 200 draws, shared across methods) and averaged their CRV, 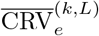. The CRV reduction at reach *L* is

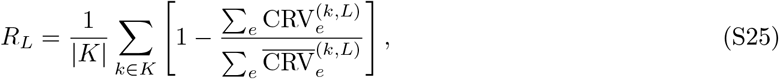

averaged over densities *d* ∈ {0.1, 0.2, 0.5, 1, 2, 5, 10} %, and we report *R*_1_, *R*_2_, *R*_3_ and their mean. We chose this density range because GRNs are generally considered sparse. *R* = 0 means the real perturbed genes explain the held-out DEGs no better than random ones, and *R* = 1 means every DEG is reached.

#### CRV reduction of graphs of known quality

For each of the ten knockdown datasets, we scored the true graph *A* (self-loops removed) and corrupted versions on all experiments with DEGs (70–76 per network), using the cells at *t* = 165–180. With *E* true edges, the corruptions were: Gaussian noise *A* + *σ N*(0, 1) on every gene pair (*σ* = 0.5, 1, 2); 0.25*E* or 0.75*E* random false edges added; true edges removed; true edges reversed; shortcut edges *i* → *k* added where *A* has *i* →*j* →*k* but no *i* → *k*; 5, 10 or 15% of all gene pairs re-drawn from Bernoulli(*ρ*) or Bernoulli(0.5), with *ρ* the true edge density; and random graphs with every pair drawn from Bernoulli(*ρ*) or Bernoulli(0.5). Each random setting was drawn ten times and averaged within each network.

#### CRV reduction by genetic-interaction type

*Norman et al*. We grouped held-out pairs using the genetic-interaction annotations from Norman et al., provided with the PerturbNet data in the GI_TYPE column of the cell metadata in `Norman_count.h5ad` (https://huggingface.co/cyclopeta/PerturbNet_reproduce/tree/main/data_paper). Of the 131 pairs, 86 were assigned to one of eight interaction types. The remaining 45 pairs, labeled “None Type”. For each type, CRV reduction was pooled over its held-out pairs as in the main analysis.

##### CaRPool-seq

Because no genetic-interaction annotations have been published for CaRPool-seq, we assigned approximate classes using features derived from the linear model of Norman et al. For each pair *AB*, we defined *G*_*AB*_ as the union of DEGs for *A, B*, and *AB*, excluding the target genes. For each perturbation group *x* ∈ {*A, B, AB*}, we calculated the mean spliced expression after library-size normalization and log(1 + *x*) transformation. We then defined *δ*_*x*_ as the difference between this mean and the control mean, restricted to *G*_*AB*_.

We fitted the following model by least squares:

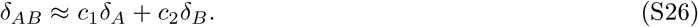

Each pair was described by the *R*^2^ of the fit and three additional features:

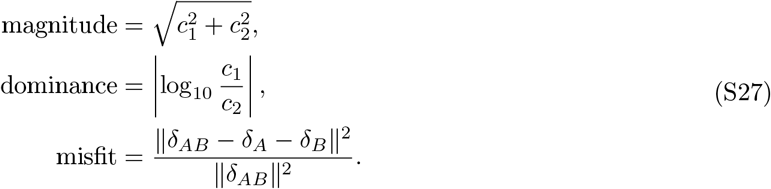

Misfit measures the deviation from the additive prediction *δ*_*A*_ + *δ*_*B*_.

We calculated the same features for the Norman et al. pairs and combined their eight annotated interaction types into four classes:

- *Additive*: approximately additive (15 pairs).
- *Synergy* : strong synergy with similar or dissimilar phenotype, and potentiation (30 pairs).
- *Dominant* : epistasis and redundant (16 pairs).
- *Poor fit* : neomorphic and suppressors (24 pairs).

One neomorphic pair, *CBARP* +*TGFBR2*, could not be matched between the annotation and feature tables and was excluded, leaving 85 pairs.

Using these 85 pairs, we trained a decision tree with a maximum depth of 3 and at least four pairs per leaf (**Supplementary Fig. S5**). The input features were misfit, ln(magnitude), dominance, and *R*^2^. Stratified five-fold cross-validation yielded an accuracy of 0.74 for the four-class classification and 0.88 for additive versus non-additive classification. We then trained the tree on all 85 pairs and applied it to CaRPool-seq. One CaRPool-seq pair, *MYB* +*RING1*, could not be scored and was left unclassified. The resulting classes approximate the Norman et al. categories and annotated by the decision tree model.

##### Feature reproducibility

To assess feature stability at the available cell numbers, we randomly split the cells in each group (control, single perturbations, and paired perturbations) into two halves and recomputed the features separately in each half. For each feature, we calculated the Spearman correlation across pairs between the two halves. We repeated this procedure for 20 random splits in each dataset.

#### Causal co-reach and genetic interactions

We asked whether the graph inferred without a perturbation pair predicts whether the two genes interact genetically. For each held-out pair (*A, B*) of Norman et al., we used the benchmark 5-fold graph in which the pair was held out, so the pair’s own cells were never used, while both single perturbations were in training. We compared non-additive pairs (any class other than approximately additive; 71 pairs) with approximately additive pairs (15 pairs), and left out pairs without a class.

Edge scores were converted to edge probabilities *p*_*e*_ exactly as for CRV: at each density of the CRV grid (0.1–10% of genes per perturbed regulator), the scores of the perturbed regulators were ranked and mapped to probabilities with the same expected number of targets for every method. The reach of gene *g* from regulator *A* is the strongest path of at most three steps,

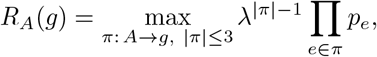

where paths run through perturbed genes (the only genes with outgoing edges), *λ* = 0.5 discounts each additional step, and *R*_*A*_(*A*) = 0. If no path exists, we set *R*_*A*_(*g*) = 0. The causal co-reach of the pair is the soft (probabilistic) Jaccard overlap of the two reach profiles:

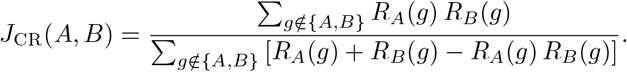

The two perturbed genes themselves are excluded. We set *J*_CR_ = 0 when the denominator is zero. Otherwise, the score is 0 when the two genes reach no common downstream genes and 1 only when they reach the same nonempty set of downstream genes, each with reach probability 1. Identical reach profiles with intermediate probabilities therefore give a score below 1. We computed *J*_CR_ at each density and averaged over the seven densities, as for CRV reduction.

We used AUROC and AUPRC to assess how well co-reach distinguishes non-additive from approximately additive pairs, and a two-sided Mann–Whitney test to compare co-reach between the two groups. For AUROC, we used co-reach as the score and non-additive pairs as positives. For AUPRC, we instead used −co-reach as the score and approximately additive pairs as positives, because additive pairs are expected to share fewer downstream genes. This choice avoids the high chance-level AUPRC of 0.83 resulting from the predominance of non-additive pairs (71 of 86); the chance level for approximately additive pairs is 0.17. We repeated this analysis using the graphs inferred in each fold by each method.

#### External support of top-ranked edges

We obtained directed regulator-to-target evidence from three sources. ChIP-Atlas target-gene tables were accessed on 22 October 2026, using hg38 and peaks within ±5 kb of the transcription start site. Edges were scored by the mean MACS2 score across experiments from the matching cell line: THP-1 for CaRPool-seq and K562 for Norman et al. CollecTRI was retrieved on 22 October 2026, and edges were scored by curation effort. The STRING v12.5 directed regulatory network (`9606.protein.regulatory.links.detailed.v12.5`) was scored by the combined score. We defined six evidence tiers (T1–T6) with increasingly strict thresholds for all three sources (**Supplementary Table S4**). An edge was evaluable if at least one source covered both its regulator and target, and supported if its evidence score met the threshold in at least one source.

For each regulator, we ranked its evaluable targets by the inferred graph edge score, excluding the regulator itself, and selected the top 2%, 5%, 10%, or 15% of edges. Edges tied at the cutoff received equal fractional selection weights. We calculated the fraction of selected edges supported by external evidence, accounting for these weights. As a random baseline, we used the supported fraction among all evaluable edges for that regulator. We averaged each fraction across regulators and defined enrichment as the ratio of the averaged selected-edge fraction to the averaged baseline fraction. For example, an enrichment of 2 indicates twice the support expected under random selection, whereas 1 indicates support equal to the random baseline. We calculated enrichment separately for each selection fraction and fold graph and reported the mean and standard deviation across the five folds.

### S4.1 Benchmarked methods and baselines

All methods received the same training and validation cells for each split. For causal discovery methods, we followed the authors’ default training procedures and hyperparameters, with adjustments for training failures or computational constraints as described below. Edge scores were extracted before any method-specific thresholding.

#### DEG baselines

These baselines construct graphs directly from DEG labels. *DEG singles* assigns an edge *j* → *g* when gene *g* is a DEG of the single perturbation of gene *j. DEG union* also assigns the DEGs of each training combination to all its targets, with weight 1*/*|*T*_*e*_| per target. This distributes the combined response equally among the perturbed genes. DEG singles was not used in the simulations because all simulated perturbations targeted multiple genes.

#### DCD-FG

DCD-FG learns a directed acyclic graph by factorizing gene–gene connections through a small set of latent factors. It uses neural networks to model conditional distributions and an augmented Lagrangian procedure to enforce acyclicity. Intervention targets are excluded from the likelihood.

We used the repository defaults for the low-rank MLP model: *m* = 20 factors, two layers with 16 hidden units each, RMSprop with a learning rate of 10^−3^, a regularization coefficient of 0.1, batch size 128, and the exponential acyclicity constraint. Training ran for up to 600 epochs using the repository’s standard stopping procedure:. Following the repository preprocessing, counts were normalized to 100,000 per cell and transformed using log(1 +*x*). Edge scores were obtained by multiplying the gene-to-factor and factor-to-gene matrices.

#### SDCD

SDCD learns a directed acyclic graph in two stages. The first stage selects candidate parents for each gene using a sparsity penalty. The second learns connections among these candidates under a spectral acyclicity penalty. Intervention targets are excluded from the likelihood.

We used the default model with 2,000 epochs per stage and early stopping. Counts were normalized to 10,000 per cell, transformed using log(1 + *x*), and z-scored. The only adjustment was the stage-1 learning rate. The default value of 2 × 10^−3^ caused the stage-1 loss to diverge on every dataset, leaving almost no candidate edges. We reduced it to 2 × 10^−4^. Edge scores were taken from SDCD’s continuous adjacency matrix, computed from the norms of the first-layer weights.

#### BICYCLE

BICYCLE models each cell’s latent expression using the steady-state distribution of a linear stochastic dynamical system. Its interaction matrix represents a gene regulatory graph that allows cycles. Interventions replace the parameters of known target genes, and a multinomial likelihood links observed counts to the latent state.

We used the authors’ large-scale Norman configuration, with a rank-5 factorized interaction matrix, an amortized encoder, and a multinomial likelihood on raw counts. We used perfect interventions with the dCas9 intervention model, RMSprop with batch size 32, and gradient clipping at 10^−3^. The loss weights were 1 for L1, 1 for KL, 0.1 for Lyapunov, and 0 for spectral regularization. Training consisted of likelihood-only pretraining followed by joint training.

The authors’ learning rate of 10^−3^ caused training to diverge on several datasets during pretraining or joint training. We tested different learning rates and pretraining durations on simulated data and selected a learning rate of 10^−4^ with 300 pretraining epochs because this combination allowed training to converge.Joint training ran for up to 1,000 epochs, with early stopping on the validation loss and patience of 50 epochs.

For Norman, a full-rank noise covariance matrix over 2,000 genes required about one hour per pretraining epoch. We therefore used a rank-15 covariance factor, matching the authors’ setting for their Perturb-seq data. We also limited pretraining to 40 hours and joint training to 110 hours. These limits ended the two stages after about 50 and 45 epochs, respectively, before early stopping was triggered.

The output edge scores were the absolute interaction weights |*β*| from the joint-training epoch with the lowest validation loss.

### S5 Other analyses

#### S5.1 Recovery of kinetic rates

For each of the 20 simulated datasets, the learned *γ*_*i*_ and *β*_*i*_*/γ*_*i*_ of the final model were compared with the simulator’s values for all 222 genes. Learned rates are identifiable only up to a common time scale per fitted model, so they were divided by their median over the genes of each dataset before pooling, and the Spearman correlation was computed over all pooled genes.

#### S5.2 Latent activity and simulated protein abundance

True protein and mRNA time courses were extracted from the simulator for the Branching and sp99 networks under knockdown and activation, expressed as log_2_ ((*x*_pert_ +0.5)*/*(*x*_ctrl_ +0.5)) of paired perturbed and control trajectories at 30 times (*t* = 105, 110, …, 250). For each experiment, 100 of its cells with their measured perturbation strengths were integrated from the control state with the trained model, mapping simulation time to the model’s latent interval so that the perturbation (*t* = 100) maps to its start and the midpoint of the training window (*t* = 172.5) to its end. Activities were averaged over the cells and divided by the activity of the model’s own unperturbed cell integrated for the same time, because the control state is not an exact steady state of the fitted model. For each perturbed regulator whose true protein changed by |log_2_ FC| ≥ 0.3 in the training window, we computed the Pearson correlation over the 30 times between its log_2_ activity and its true protein curve.

### S5.3 CD11b analysis

CausalODE was trained on all CaRPool-seq GMP cells, holding out 10% of the pairs (19 of 158) only for checkpoint selection. For each single knockdown with at least 20 GMP cells, we report the target’s normalized spliced RNA relative to control cells, the mean latent activity of the targeted regulator at the end of integration relative to the model’s unperturbed cell, and the edge probability from the regulator to *ITGAM*. CD11b surface protein was taken from the antibody counts, normalized to 10,000 counts per cell; its log_2_ fold change is the group mean over the control mean, tested per cell on log(1 + *x*) values with a two-sided Mann–Whitney test against control cells and Benjamini–Hochberg corrected across the 28 knockdowns. Target-RNA differential expression is from the DEG analysis above.

### S6 Supplementary Figures

**Figure S1:**
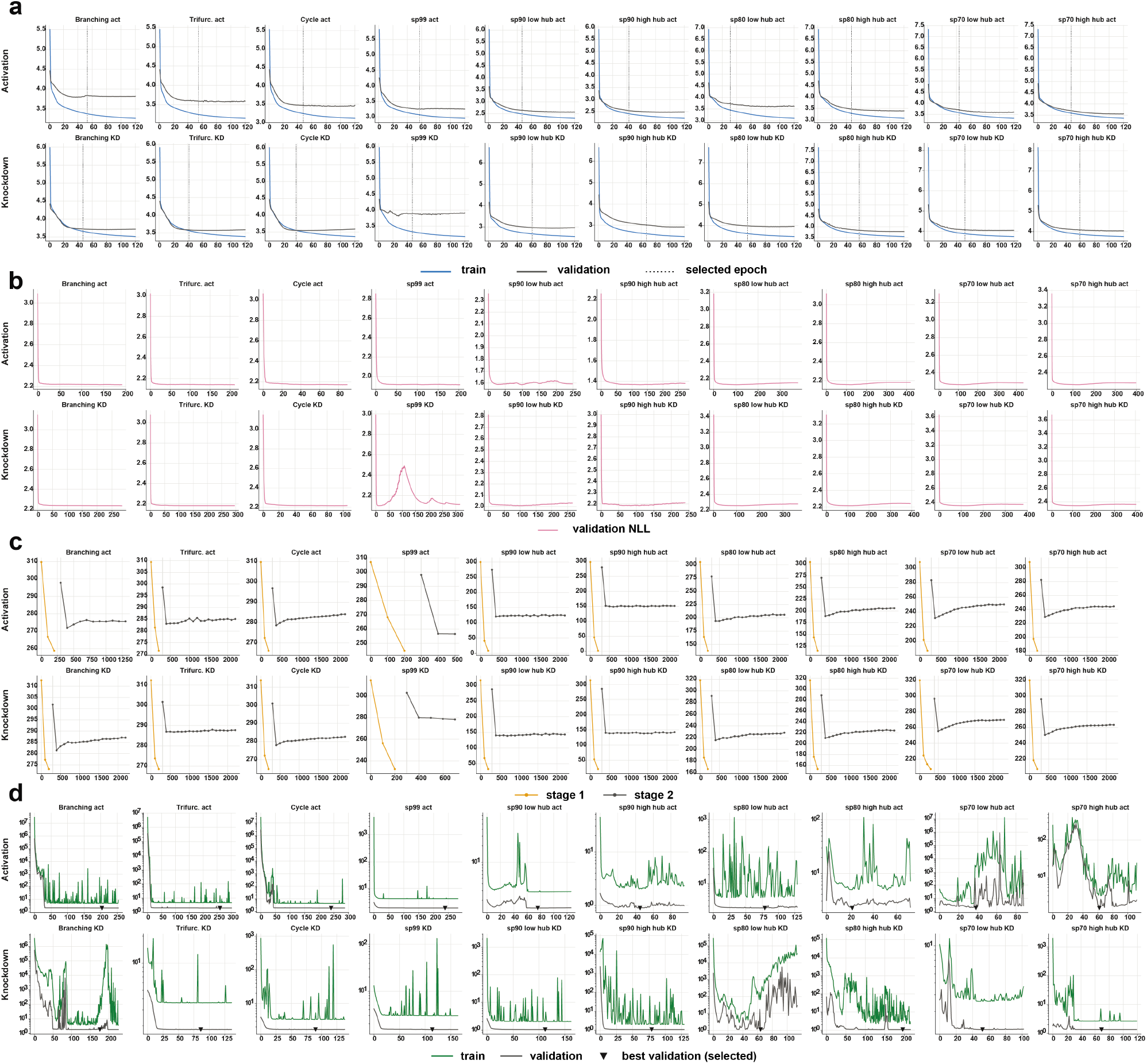
Training curves of benchmarked methods on simulated datasets. Within each panel, columns show the ten simulated networks, and rows show activation (top) and knockdown (bottom). For panels (b)–(d), each method uses its default early-stopping rule. **(a)** CausalODE training (blue) and validation (grey) total loss. Dotted lines mark selected checkpoints. **(b)** DCD-FG validation negative log-likelihood. **(c)** SDCD training objectives for stage 1 (orange) and stage 2 (grey), logged every 100 epochs. Stage 2 follows stage 1 on the epoch axis. **(d)** BICYCLE joint-training loss on a log scale: training (green) and validation (grey). Triangles mark checkpoints with the lowest validation loss.

**Figure S2:**
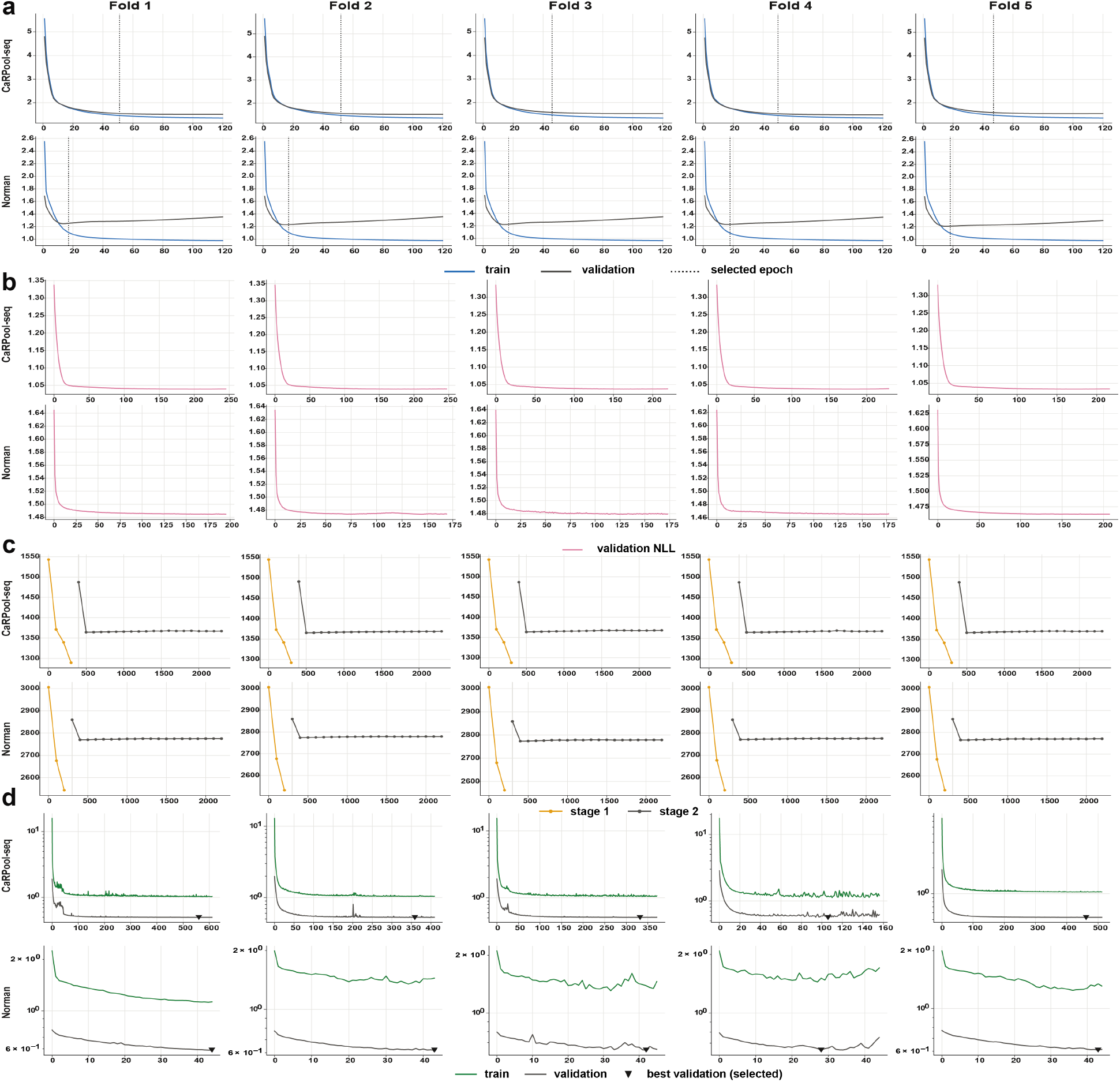
Training curves of benchmarked methods on real Perturb-seq datasets. Within each panel, columns show the five train–test splits, and rows show CaRPool-seq (top) and Norman et al. (bottom). Stopping rules and runtime limits follow the benchmark settings described in the Methods. **(a)** CausalODE training (blue) and validation (grey) total loss. Dotted lines mark selected checkpoints. **(b)** DCD-FG validation negative log-likelihood. **(c)** SDCD training objectives for stage 1 (orange) and stage 2 (grey). Stage 2 follows stage 1 on the epoch axis. **(d)** BICYCLE joint-training loss on a log scale: training (green) and validation (grey). Triangles mark checkpoints with the lowest validation loss.

**Figure S3:**
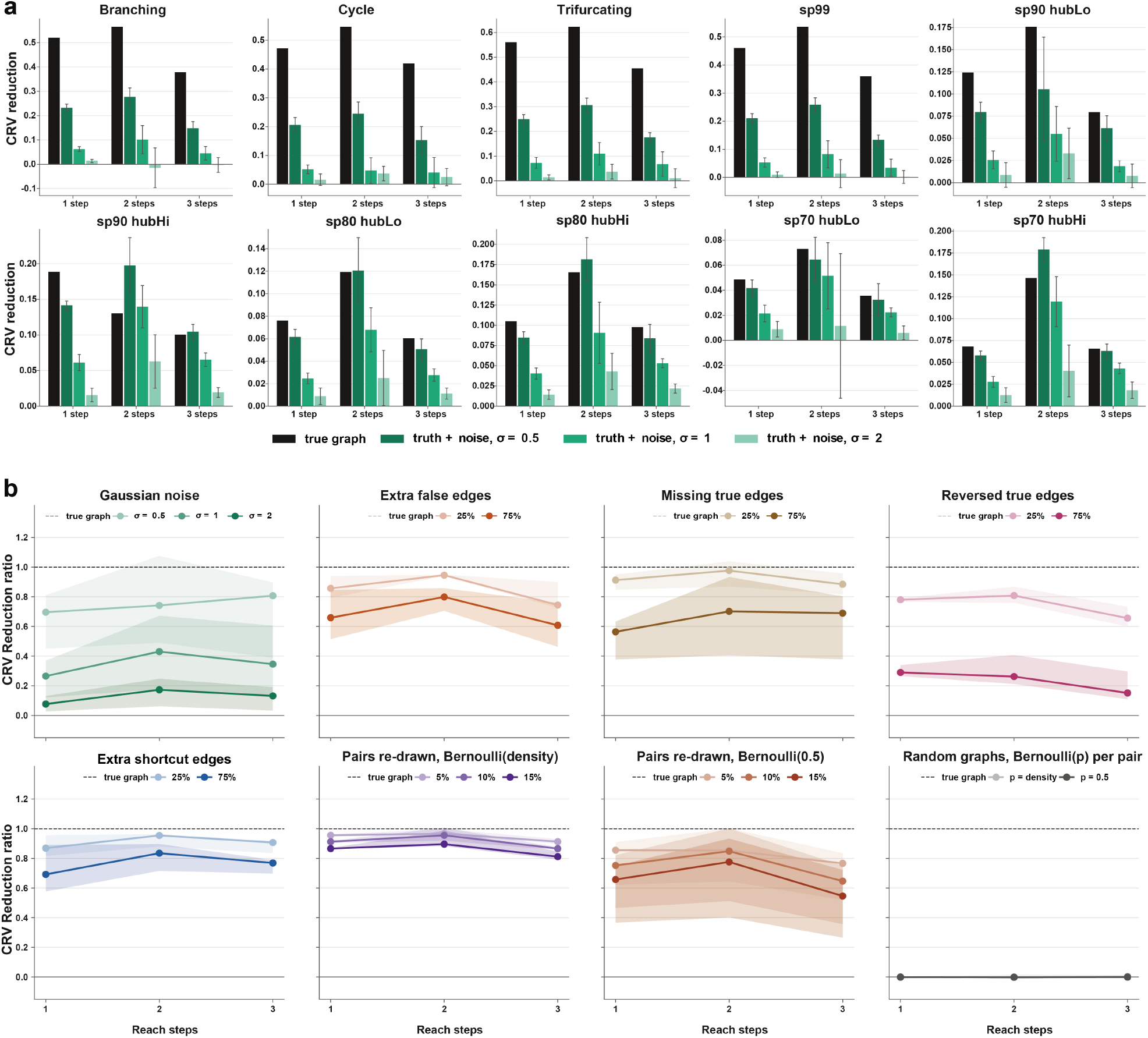
Sensitivity of CRV reduction to graph errors. Scores are evaluated on all experiments with DEGs in ten simulated knockdown datasets, using one-, two-, and three-step reach and the same scoring procedure as the real-data benchmark. Self-edges are excluded. **(a)** CRV reduction for true graphs (black) and graphs with Gaussian noise *N*(0, *σ*^2^) added to every gene pair (*σ* = 0.5, 1, and 2; green, dark to light). Noisy-graph bars and error bars show the mean and standard deviation over ten draws. **(b)** CRV reduction relative to the true graph at the same path length (dashed line, 1). Errors include Gaussian noise, added false edges, missing edges, reversed edges, added shortcuts, and randomly resampled gene pairs. Shortcuts connect genes linked by a two-step path without a direct edge. Edge modifications affect 25% or 75% of the true edge count; pair resampling affects 5%, 10%, or 15% of all gene pairs. Resampled pairs and random graphs use Bernoulli probabilities equal to the true graph density or 0.5. Random modifications are averaged over ten draws per network. Lines and shading show medians and interquartile ranges across networks.

**Figure S4:**
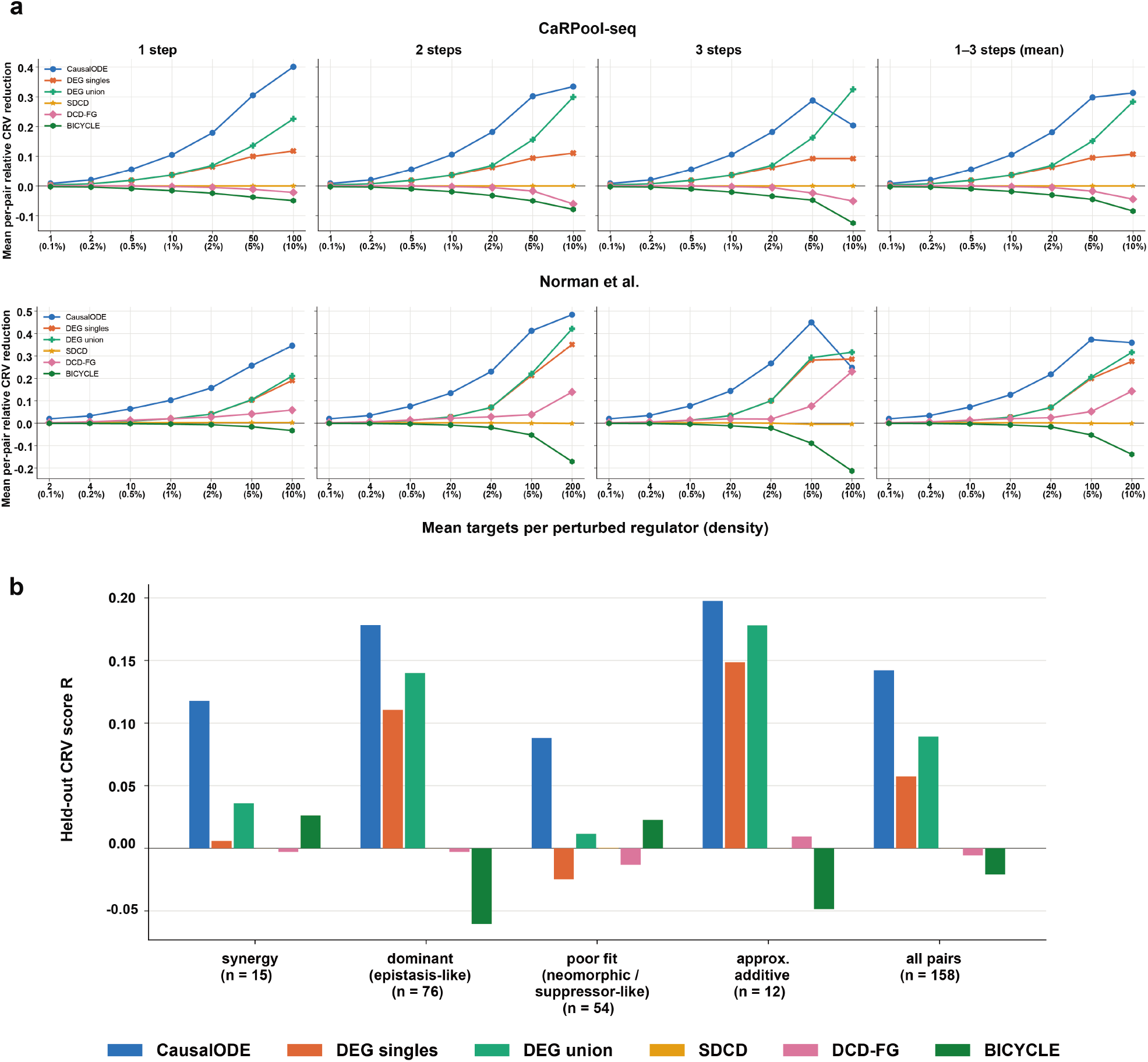
CRV reduction across graph densities and interaction classes. **(a)** Mean per-pair CRV reduction across five train–test splits for CaRPool-seq (158 pairs, top) and Norman et al. (131 pairs, bottom). Columns show one-, two-, and three-step reach and their mean. The x-axis shows the expected number of targets per regulator, with graph density in parentheses. Methods are evaluated at matched densities. **(b)** One-step CRV reduction for held-out CaRPool-seq pairs, pooled within each interaction class. In the absence of published annotations, classes were assigned using a depth-3 decision tree trained on Norman et al. interaction metrics (five-fold cross-validated accuracy, 0.74). The four classes are synergy, dominant, poor fit, and approximately additive. These labels are approximate because the interaction metrics are less reproducible in CaRPool-seq than in Norman et al.

**Figure S5:**
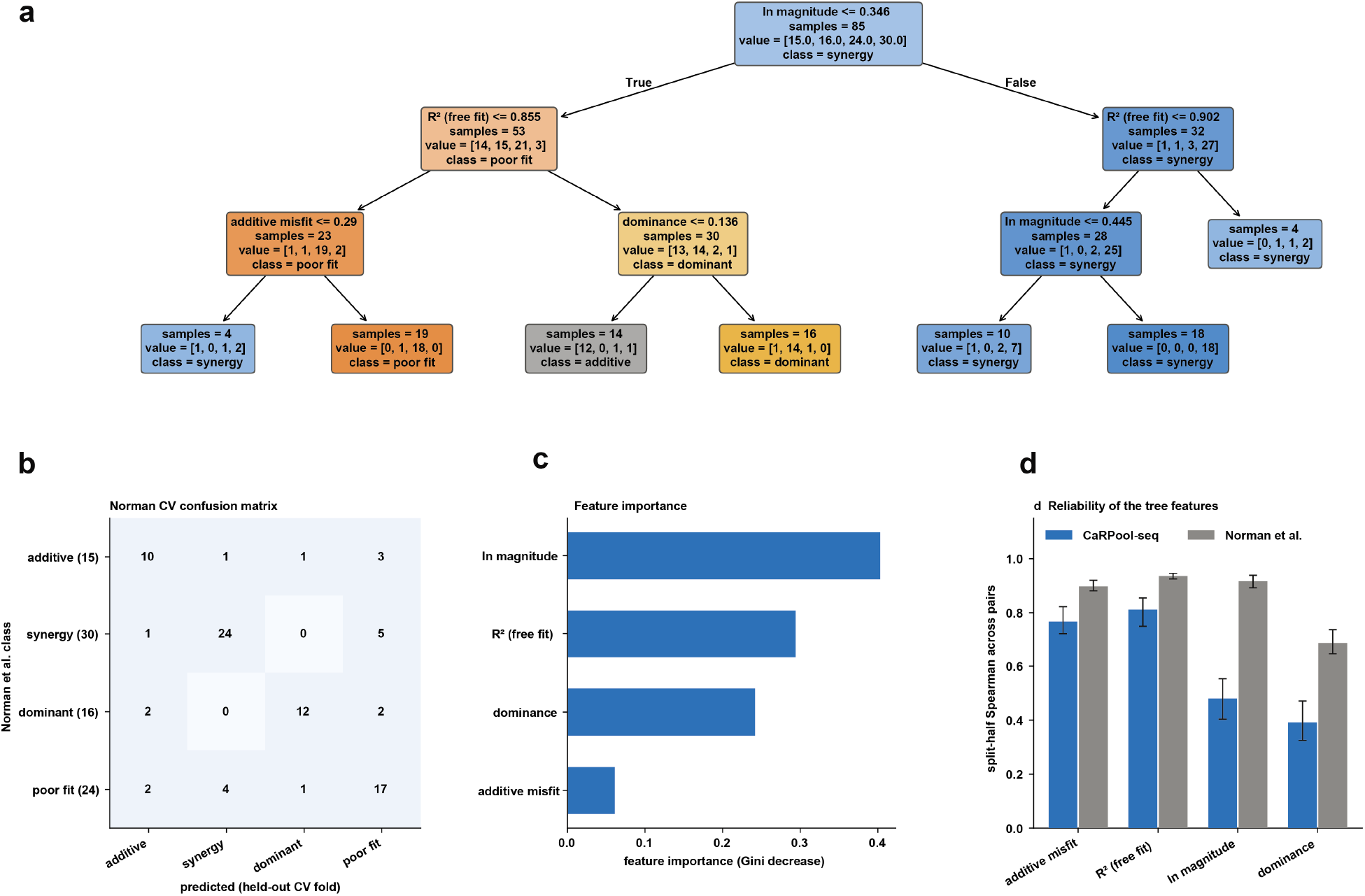
Assigning genetic-interaction classes to CaRPool-seq pairs. Interaction features were derived from the linear model *δ*_*AB*_ = *c*_1_*δ*_*A*_ + *c*_2_*δ*_*B*_, where *δ* denotes expression changes relative to controls. Features describe effect magnitude, dominance, model fit, and deviation from additivity. **(a)** A depth-3 decision tree trained on 85 annotated Norman et al. pairs, grouped into four classes: additive (approximately additive), synergy (strong synergy and potentiation), dominant (epistasis and redundant), and poor fit (neomorphic and suppressor). Node counts follow the order additive, dominant, poor fit, and synergy. This tree assigned the CaRPool-seq interaction classes used in Supplementary Fig. S4b. **(b)** Stratified five-fold cross-validation confusion matrix, with accuracy of 0.74 for the four classes and 0.88 for additive versus non-additive pairs. **(c)** Feature importance measured by the decrease in Gini impurity. **(d)** Split-half reproducibility of the interaction features across 20 random cell splits. Bars show median Spearman correlations; error bars show the 10th–90th percentiles. Lower reproducibility in CaRPool-seq indicates that its assigned classes should be treated as approximate.

**Figure S6:**
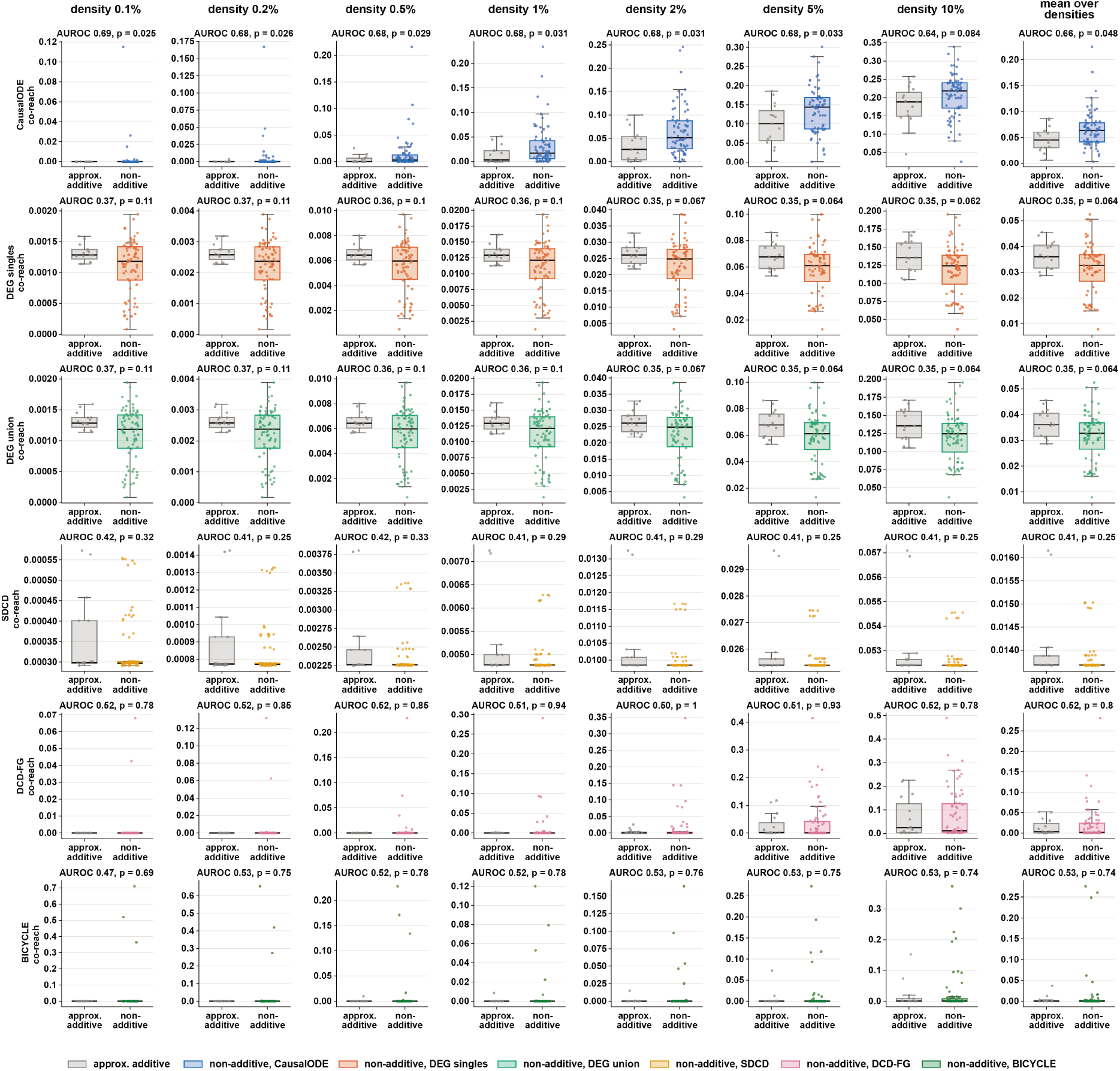
Causal co-reach by genetic-interaction type. Causal co-reach of approximately additive (grey, *n* = 15) and non-additive (method colour, *n* = 71) Norman et al. pairs, computed using graphs trained without the corresponding pairs. Rows show methods; columns show the seven graph densities and their mean. Boxes show medians and interquartile ranges; whiskers extend to 1.5 times the interquartile range; points represent individual pairs. Panel titles report AUROC with non-additive pairs as positives and unadjusted two-sided Mann–Whitney *p*-values.

**Figure S7:**
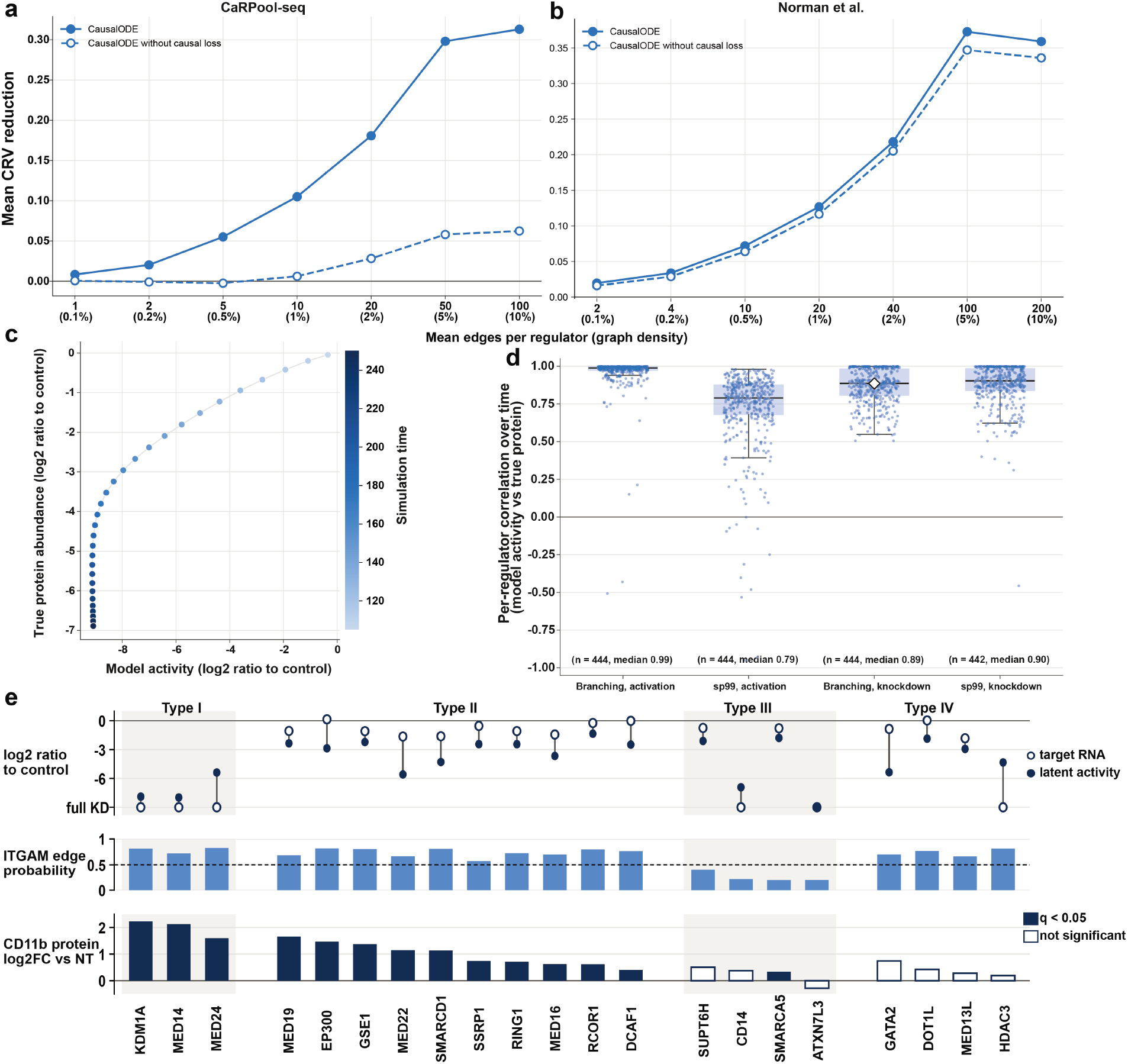
Causal-loss ablation and latent mediator activity. **(a, b)** Mean held-out CRV reduction for CaRPool-seq (a) and Norman et al. (b), averaged over one–three reach steps. Solid/dashed lines: with/without the causal loss. **(c)** Latent activity versus true protein abundance for a knocked-down regulator in the Branching simulation at 30 time points, expressed as log_2_ ratios to their respective time-matched un-perturbed controls (Pearson *r* = 0.89). **(d)** Temporal Pearson correlations computed as in (c) for Branching and sp99 under activation and knockdown. Points represent experiment–regulator pairs with true protein changes of |log_2_ FC| ≥ 0.3 (*n* = 442–444 per dataset). Boxes show medians and quartiles; the diamond marks (c). **(e)** Top: target RNA and latent activity relative to controls (log_2_); “full KD” indicates un-detected RNA. Middle: regulator-to-*ITGAM* edge probability. Bottom: CD11b protein log_2_ fold change versus non-targeting controls; filled bars indicate significance (Mann–Whitney test, BH correction across 28 knockdowns, *q <* 0.05). Types I/II: weaker/stronger activity reductions than RNA reductions, with high edge probabilities and increased CD11b. Type III: low edge probabilities and small CD11b changes. Type IV: high edge probabilities without significant CD11b changes.

**Figure S8:**
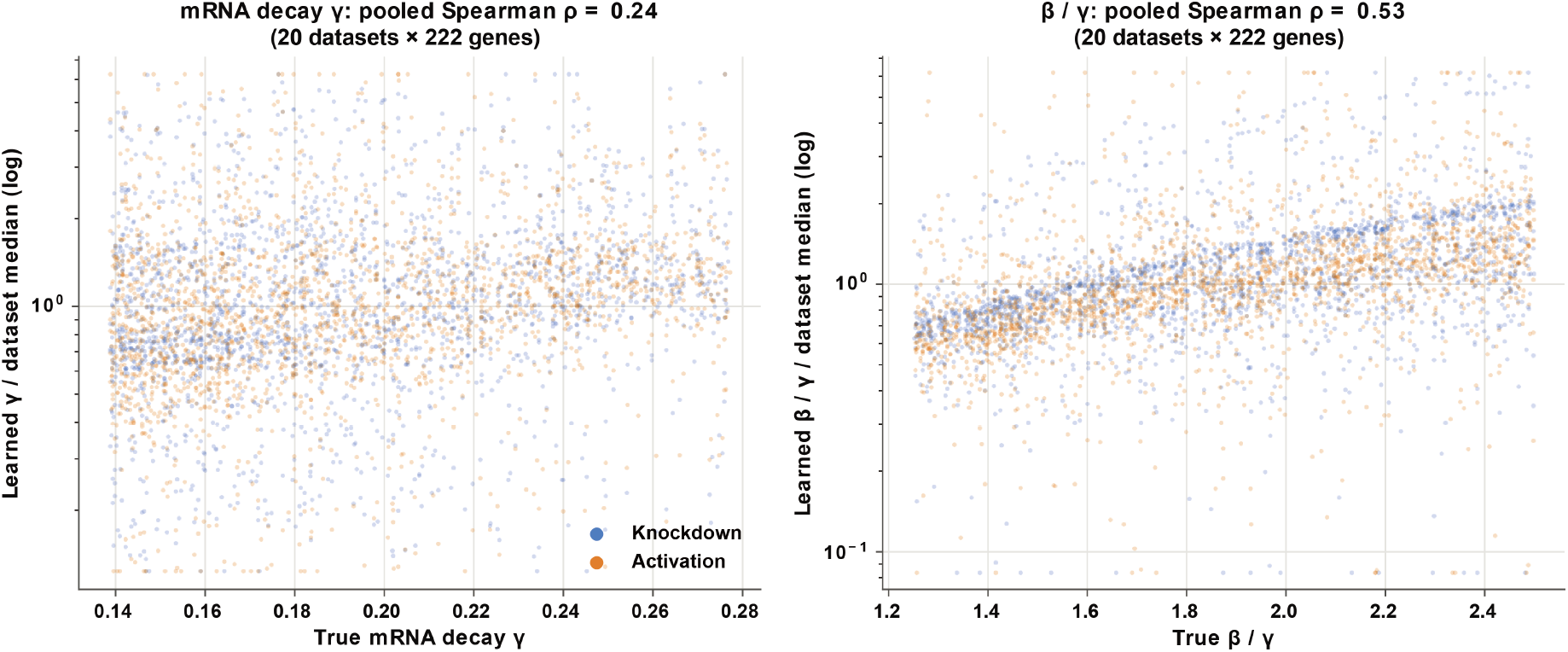
Recovery of RNA kinetic rates in simulations. Learned versus true mRNA degradation rates *γ* (left) and splicing-to-degradation ratios *β/γ* (right) across 20 datasets (4,440 observations in total). Blue and orange indicate knockdown and activation, respectively. The simulator uses the same splicing rate for all genes, so *β* is evaluated through its ratio to *γ*. Learned values were divided by their within-dataset median before pooling; true values were unchanged. Panel titles report pooled Spearman correlations: 0.24 for *γ* and 0.53 for *β/γ*.

**Figure S9:**
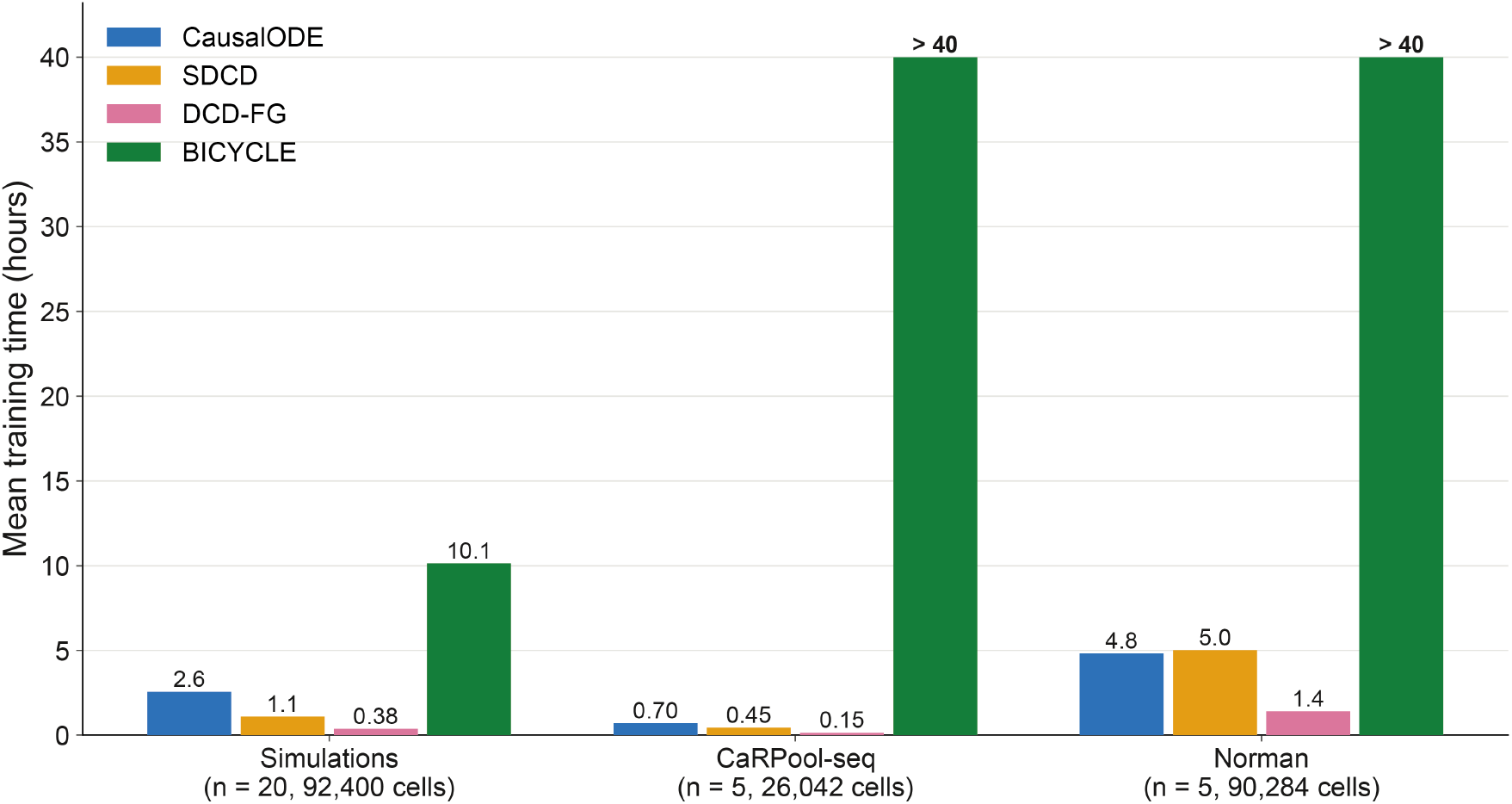
Training time of graph-inference methods. Mean wall-clock time per model using one GPU per run, excluding data loading and preprocessing. Results cover 20 simulated datasets and five folds each of CaRPool-seq and Norman et al. Labels indicate the number of datasets or folds (*n*) and cells per dataset before splitting. BICYCLE averaged 51.8 hours on CaRPool-seq (range, 30–72 hours) and reached the 110-hour time limit on every Norman et al. fold.

### S7 Supplementary Tables

**Table S1:** Simulated benchmark graphs (222 genes). Edges column excludes self-loops and feedback edges column is the percentage of edges that lie on a directed cycle.

| Graph | Edges | Sparsity<br>(%) | Self-loops | Repressive<br>(%) | Max out<br>degree | Max in<br>degree | Feedback<br>edges (%) |
|---|---|---|---|---|---|---|---|
| Branching | 452 | 99.1 | 20 | 25 | 18 | 6 | 15 |
| Trifurcating | 341 | 99.3 | 12 | 25 | 12 | 3 | 7 |
| Cycle | 361 | 99.3 | 22 | 21 | 6 | 3 | 47 |
| sp99 | 453 | 99.1 | 20 | 23 | 21 | 7 | 42 |
| sp90, low hub | 4,908 | 90.0 | 20 | 13 | 97 | 40 | 73 |
| sp90, high hub | 4,908 | 90.0 | 20 | 13 | 178 | 31 | 74 |
| sp80, low hub | 9,837 | 80.0 | 20 | 7 | 102 | 57 | 73 |
| sp80, high hub | 9,837 | 80.0 | 20 | 7 | 179 | 56 | 74 |
| sp70, low hub | 14,765 | 69.9 | 20 | 4 | 104 | 85 | 74 |
| sp70, high hub | 14,765 | 69.9 | 20 | 4 | 180 | 82 | 74 |

**Table S2:** CRV reduction on held-out perturbation pairs pooled across five folds (CaRPool-seq, 158 pairs; Norman et al., 131 pairs). The 1–3-step score is the mean across all three path lengths. Higher scores indicate better graph quality. The highest value in each column is bold.

|  | CaRPool-seq |  |  |  | Norman et al. |  |  |  |
|---|---|---|---|---|---|---|---|---|
| Method | 1 step | 2 steps | 3 steps | 1–3 steps | 1 step | 2 steps | 3 steps | 1–3 steps |
| CausalODE | **0.155** | **0.146** | **0.125** | **0.142** | **0.140** | **0.200** | **0.178** | **0.173** |
| DEG union | 0.077 | 0.093 | 0.097 | 0.089 | 0.057 | 0.118 | 0.114 | 0.096 |
| DEG singles | 0.057 | 0.058 | 0.057 | 0.057 | 0.054 | 0.108 | 0.115 | 0.093 |
| SDCD | 0.000 | 0.000 | 0.000 | 0.000 | 0.003 | 0.001 | 0.000 | 0.001 |
| DCD-FG | −0.004 | −0.007 | −0.006 | −0.006 | 0.025 | 0.041 | 0.054 | 0.040 |
| BICYCLE | −0.018 | −0.022 | −0.023 | −0.021 | −0.009 | −0.032 | −0.040 | −0.027 |

**Table S3:** Candidate-matrix coverage of the external evidence sources. Coverage is the number of regulator–target pairs that can be evaluated by a source, divided by the complete off-diagonal regulator-by-gene candidate matrix. The Cas13 matrix contains 28 × 999 = 27,972 pairs, and the Norman matrix contains 105 × 1999 = 209,895 pairs. The combined reference uses the union of matched-cell-type ChIP-Atlas, CollecTRI, and STRING regulatory coverage.

| Dataset | Evidence source | Evaluable pairs | Matrix coverage |
|---|---|---|---|
| Cas13 | Matched ChIP-Atlas | 5,994 | 21.4% |
|  | CollecTRI | 8,991 | 32.1% |
|  | STRING regulatory | 19,752 | 70.6% |
|  | **Combined, any source** | **22,511** | **80.5%** |
| Norman | Matched ChIP-Atlas | 29,985 | 14.3% |
|  | CollecTRI | 69,965 | 33.3% |
|  | STRING regulatory | 105,280 | 50.2% |
|  | **Combined, any source** | **122,022** | **58.1%** |

**Table S4:** Confidence tiers for the combined external-support analysis. At each tier, an edge was labeled as externally supported if at least one available source met its corresponding threshold. ChIP-Atlas scores were computed for matched cell types within ±5 kb of the target transcription start site. CollecTRI uses OmniPath curation effort, defined as the number of unique database–literature-reference pairs. STRING values are directed regulatory combined scores. Higher tiers impose progressively more stringent evidence thresholds.

| Tier | ChIP-Atlas score | CollecTRI effort | STRING regulatory score |
|---|---|---|---|
| T1 | 50 | 1 | 150 |
| T2 | 100 | 2 | 250 |
| T3 | 200 | 3 | 400 |
| T4 | 500 | 5 | 600 |
| T5 | 800 | 10 | 700 |
| T6 | 1,000 | 20 | 800 |

